# Top-down Sequencing of Intact Proteoforms using the timsOmni mass spectrometer: Accurate Determination of Co-occurring Histone Modifications

**DOI:** 10.64898/2026.05.01.722147

**Authors:** Francis Berthias, Nurgül Bilgin, Athanasios Smyrnakis, Elisa Le Boiteux, Mariangela Kosmopoulou, Christian Albers, Detlev Suckau, Jasmin Mecinović, Dimitris Papanastasiou, Ole N. Jensen

**Author notes:** Corresponding author: Ole N. Jensen, Department of Biochemistry and Molecular Biology, University of Southern Denmark, Campusvej 55, 5230 Odense, Denmark.

## Abstract

Deep characterization of intact proteoforms remains an analytical challenge in functional proteomics, particularly for heterogenous multi-site post-translational modifications at distinct amino acid residues. Histones are among the most dynamically and diversely post-translationally modified proteins in eukaryote cells, carrying multiple, co-occurring and reversible modifications that can give rise to isomeric proteoform species. Tandem mass spectrometry with multimodal fragmentation capabilities is a promising approach for deep characterization of intact proteoforms, such as modified histones. We applied the novel timsOmni mass spectrometer, which incorporates the Omnitrap platform enabling multimodal MSⁿ workflows, for residue-level mapping of histone modifications, including acetylation and methylation. Recombinant histones H3.1 and H4 were *in vitro* acetylated by enzymes GCN5, PCAF and p300 to generate mono- and multi-acetylated proteoforms. Complementary MS^2^ electron- and collision-based dissociation (ECD, EID, _R_CID and ECciD), together with MS^3^ strategies, produced complete or near-complete backbone fragmentation of intact protein ions (>92% amino acid sequence coverage). For monoacetylated species generated by the more site-selective lysine acetyltransferases, the dominant proteoform matched the known catalytic preferences of the enzymes (H3.1K14ac for GCN5 and PCAF, and H4K8ac for PCAF), while minor positional isomers were also identified and their relative abundance estimated. In contrast, the broader substrate specificity of p300 produced a wide distribution of H4 proteoforms bearing up to seven acetylated lysine residues. Species carrying six and seven acetylations were characterized by multimodal MS^2^/MS^3^ experiments, enabling localization of individual acetylation sites and discrimination of positional isomers. Finally, endogenous histone proteoforms from liver extracts were analyzed, yielding sequence coverages of 92–93% for the most abundant species and enabling confident localization of multiple PTMs (acetylation and methylation). These results illustrate that multimodal MS^n^ fragmentation of intact proteins supports residue-level assignment of combinatorial histone marks and coexisting positional isomers.

**Graphical Abstract:** 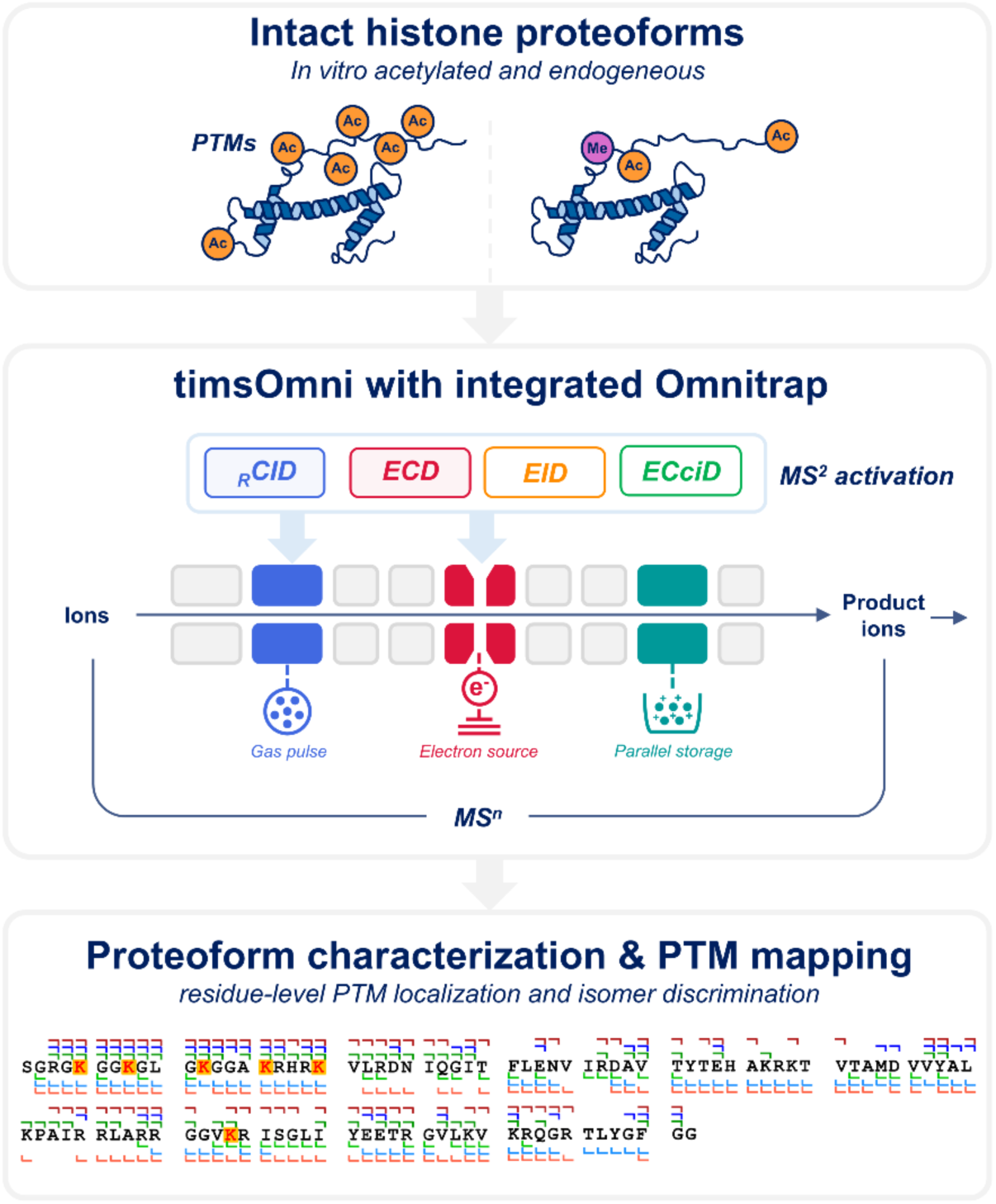

**Highlights:**

- Multimodal MS²/MS³ maps histone PTMs on intact proteins.
- ECD, EID, _R_CID, and ECciD provide complete or near-complete sequence coverage.
- MS³ localizes acetylation sites, distinguishes positional isomers.
- Endogenous H4 proteoforms are assigned with site-specific PTM mapping.

## Introduction

The diversity of the eukaryote proteome arises from alternative splicing, sequence variants, and post-translational modifications (PTMs) of proteins, that extend beyond the information encoded in the 20,000 human genes, thus generating millions of distinct proteoforms (1–3). PTMs modulate protein function by introducing chemical modifications at specific amino acid residues within proteins. Many modifications are reversible and dynamically alter or modulate protein structure, interactions, localization, and activity (4–6). The actual site of location, the chemical nature and the combination of PTMs in a protein dictate the biological outcome. To date, more than 500 discrete PTMs have been described (7–9).

Histones are among the most densely post-translationally modified proteins in cells. These small basic proteins package the DNA into nucleosomes and modulate chromatin structure and gene accessibility via site-specific and combinatorial patterns of PTMs, often referred to as the “histone code” (10, 11). More than 200 distinct histone PTM types have been reported, dominated by lysine acetylation, lysine/arginine methylation, and serine/threonine phosphorylation. The combinatorial distribution of different co-occuring PTMs across individual histones provides a vast proteoform space, exceeding that observed for many other cellular proteins.

The biological function of histone PTMs depends not only on their presence but also on their precise residue localization. Among them, lysine acetylation (Kac) represents a central regulator, as it modulates histone-DNA interactions through neutralization of the ε-amino group of lysine side chains, weakening histone-DNA electrostatic interactions and promoting recruitment of acetyl-lysine reader proteins (12). Importantly, the same modifications on a different residue can have distinct regulatory consequences. For example, acetylation of histone H3 at K9 is associated with active promoters and early transcriptional steps, while K27ac marks active enhancers and correlates with transcriptional output and elongation (13–16). Similarly, acetylation of histone H4 at K16 alters higher-order chromatin folding, in contrast to acetylation of other H4 lysine residues (K5, K8 and K12) (17). Defects in histone PTMs deposition and removal are implicated in numerous diseases including cancer (18–20), or neurodegenerative disorders (21). Consequently, interpretation of histone PTM function requires analytical strategies capable of determining both PTM identity and its precise site of occurrence.

Antibody-based assays and bottom-up proteomics by mass spectrometry (MS) are the standard approaches for identifying individual histone PTMs (22–24). However, bottom-up workflows rely on proteolytic digestion into short peptides, which inherently lose the link between distantly located PTM sites, while the antibodies assays are often limited by epitope occlusion (25, 26). Although partial PTM connectivity can be retained using middle-down proteomics strategies (27–29) or alternative proteases designed to detect local PTM crosstalk (30), preservation of the complete combinatorial pattern of histone PTMs generally requires analysis of intact histone proteoforms.

Top-down mass spectrometry analysis aims to address this limitation by directly analyzing intact proteins and preserving full proteoform information (PTM identification and localization, protein sequence variants, truncations). Nevertheless, unambiguous localization of PTMs at the protein level remains analytically challenging. First, the combinatorial space of positional PTM isomers increases rapidly with the number of potential modification sites, complicating proteoform identification when sequence coverage is incomplete, in contrast to peptide-level analyses where the number of potential modification sites is inherently limited. Second, co-isolation of positional PTM isomers can generate chimeric mass spectra, complicating unambiguous proteoform assignment. Finally, many PTMs introduce only small mass differences relative to the intact protein mass. For example, acetylation and trimethylation differ by only 35.4 mDa, while citrullination of arginine produces a +0.98 Da mass shift, requiring high-quality fragment ion information for confident PTM assignments. Separation strategies applied prior to MS analysis can mitigate these challenges by reducing spectral complexity. Liquid chromatography (LC) can separate distinct proteoform families at intact protein level (31) and separation by degree of acetylation or methylation has been demonstrated for histone tails in middle-down workflows (∼5 kDa) (32, 33). However, PTM positional isomers often exhibit similar retention behavior. Gas-phase separation such as ion mobility spectrometry (IMS) provides an additional dimension of separation based on ion structure (34–36) and has demonstrated high isomeric separation capabilities for large molecular ions (37–40), including targeted separation of positional isomers of acetylated histones (41, 42).

Even with effective separation, unambiguous PTM site assignment depends on the informative content of tandem mass spectra and the extent of backbone fragmentation. Collision-based dissociation (e.g., CID or HCD) typically yields limited sequence coverage and is biased toward preferential cleavage pathways. Electron-based dissociation methods (ECD/ETD) generate more extensive backbone fragmentation and are particularly suited for large, highly charged protein ions while preserving labile PTMs (43–45). Combining complementary ion activation methods into multimodal workflows is a powerful strategy to reach near complete amino acid sequence coverage (46). MS^n^ (n≥2) and hybrid workflows can effectively reduce the theoretical proteoform search, enabling targeted interrogation of informative fragment ions (47, 48). Furthermore, recent computational evidence suggests that the addition of internal fragment ions may be critical for expanding effective sequence coverage and achieving unambiguous characterization of complex histone PTM patterns (49). Altogether, this underscores that top-down proteomics remains an actively developing analytical workflow, in which fragmentation strategies and data analysis approaches are still being refined to fully exploit the complexity of the recorded information.

In this study, we leverage the timsOmni^TM^ platform (Bruker Daltonics), which integrates the Omnitrap^®^ segmented linear ion trap enabling multiple complementary activation methods within a time-of-flight mass spectrometer. The system supports resonance collision-induced dissociation (_R_CID), electron-capture dissociation (ECD), electron-induced dissociation (EID), as well as hybrid activation strategies,, such as electron-capture collision-induced dissociation (ECciD). Using *in vitro* enzymatically acetylated histones H3.1 and H4, we exploit the timsOmni platform to localize acetylation sites, resolve positional isomers and compare complementary fragmentation pathways, including MS³ analysis. We demonstrate that multimodal MSⁿ analysis enables complete or near complete sequence coverage, allowing confident, site-resolved mapping of PTMs in intact histone proteoforms isolated from human cells.

## Experimental Procedures

### Histone proteins and in vitro enzymatic acetylation

Recombinant unmodified histone H4 (*Xenopus laevis* sequence, monoisotopic neutral mass of 11,229.4 Da) was expressed and purified in-house as descrived previously (50). The *Xenopus laevis* H4 amino acid sequence is identical to the canonical human histone H4 sequence. Recombinant human PCAF and GCN5 were expressed and purified in-house as described previously (51). Human p300 was purchased from Active Motif and used without further purification. Recombinant human histone H3.1 (monoisotopic neutral mass of 15,263.4) was purchased from Sigma-Aldrich.

Enzymatic acetylation reactions were performed in vitro in a total volume of 30 µL using a reaction buffer composed of 50 mM HEPES, 0.1 mM EDTA, and 1 mM DTT at pH 8.0. Acetyl-coenzyme A (acetyl-CoA, Sigma-Aldrich) was used as co-substrate at a final concentration of 100 µM. Histone proteins were added to a final concentration of 20 µM. Recombinant lysine acetyltransferases (p300, PCAF, or GCN5) were diluted in reaction buffer and added to initiate the reaction, yielding a final enzyme concentration of 100 nM (enzyme-to-substrate molar ratio ∼1:200). Reactions were carried out at 37 °C with agitation (750 rpm) using a thermomixer (Eppendorf) for 10 min, 30 min and 60 min for GCN5, PCAF and p300 reactions, respectively.

Following incubation, enzyme reactions were quenched by addition of 10% (v/v) trifluoroacetic acid (TFA) or formic acid (FA). Samples were subsequently prepared for mass spectrometry by buffer exchange using centrifugal ultrafiltration units equipped with polyethersulfone (PES) membranes (Vivaspin, Sartorius). A 3 kDa MWCO filter (VS0191) was used for histone H4, and a 5 kDa MWCO filter (VS0111) for histone H3.1. Membranes were pre-conditioned with Milli-Q water prior to sample loading. Quenched reaction mixtures were diluted up to ten-fold with Milli-Q water, centrifuged (5,000–8,000g), and washed at least twice to remove non-volatile buffer components while restoring the retentate volume to approximately the initial volume. Protein concentration after buffer exchange was determined using a NanoPhotometer N60 (Implen).

### Endogeneous histone preparation

Histones were extracted from frozen cell pellets (human hepatocellular carcinoma cell line HepG2/C3A) by acid extraction (adapted from Garcia et al., 2008) (52). Briefly, cell pellets were resuspended in nucleus isolation buffer (15 mM Tris-HCl pH 7.5, 60 mM KCl, 11 mM CaCl_2_, 5 mM NaCl, 5 mM MgCl_2_, 250 mM sucrose, 1 mM dithiothreitol, 10 mM sodium butyrate and 0.1% Igepal) supplemented with protease inhibitors (cOmplete™ Protease Inhibitor Cocktail, Roche) and phosphatase inhibitors (PhosSTOP, Roche). Nuclei were isolated by centrifugation (1,000g, 5 min) and washed twice with nucleus isolation buffer without Igepal. Histones were extracted by resuspending the pellet in 0.2 M H_2_SO_4_ for 1 hour under gentle agitation, and the supernatant was collected after centrifugation (20,000g, 5 min). Histones were precipitated by addition of trichloroacetic acid to a final concentration of 20% and incubated overnight. Precipitated proteins were recovered by centrifugation (20,000g, 15 min) and washed once with 0.1% HCl in acetone and twice with pure acetone. Pellets were air-dried and resuspended in ultrapure H_2_O. All steps were performed at 4°C.Histone purity was assessed by SDS-PAGE and protein concentration was determined using a NanoPhotometer N60 (Implen).

Acid-extracted histones were subjected to off-line reversed-phase micropurification. Microcolumns were prepared by packing a small C18 disk and reversed-phase resin (POROS R1 Applied Biosystems) into 10 µL pipette tips. The packed material was equilibrated with 100% acetonitrile followed by 0.1% formic acid in Milli-Q water. Histone samples, acidified in 0.1% formic acid, were loaded onto the microcolumn and washed twice with 0.1% formic acid. Proteins were then eluted stepwise using increasing concentrations of acetonitrile () in 0.1% formic acid to simulate a reversed-phase LC gradient under off-line conditions. Fractions corresponding to defined organic solvent percentages were collected separately. The fraction eluting at 25-35% acetonitrile, corresponding to the analytical LC retention window of H4, was selected for subsequent analysis.

### timsOmni prototype instrument

All experiments were performed on a prototype timsOmni platform (Bruker Daltonics), generated through extensive modification of a timsTOF Ultra system, and recently described for top-down analysis of intact proteins and oligonucleotides (53, 54). The prototype retains the the timsTOF Ultra front-end, a reconfigured quadrupole mass filter, a upgraded collision cell, a high-resolution orthogonal time-of-flight (oTOF) mass analyzer, and integrates the latest version of the Omnitrap platform (48). Samples were introduced by direct infusion using a nano-electrospray ionization (nESI) source (NEOS, Bruker) with an ESI voltage of 1.0-1.4 kV. Samples were analyzed at a final concentration of 1-5 µM in water/methanol/formic acid (49:50:1). Electrosprayed ions were introduced into the mass spectrometer through a 1.0 mm inner diameter resistive glass capillary into differentially pumped regions comprising RF ion funnels and quadrupole ion guides for efficient transmission and collisional cooling. The instrument also incorporates a trapped ion mobility spectrometry (TIMS) analyzer, which was not employed in the present study and was set in transparent mode.

Precursor ions were selected using the quadrupole mass filter upstream of the Omnitrap platform. Selected ions were transferred into the Omnitrap through an RF hexapole and DC ion optics. The Omnitrap is a linear ion trap composed of nine hyperbolic quadrupole segments (Q1-Q9), with spatially separated regions for ion accumulation, isolation and ion enrichment. It provides radial confinement through variable-frequency rectangular waveforms and enables dynamic axial ion manipulation via independently applied DC potentials across each segment. A Q10 transfer section guides processed ions from the Omnitrap toward the collision cell and TOF analyzer.

Resonance CID (_R_CID) was performed in Q2 by dipolar excitation of trapped ions in the presence of a pulsed N₂ buffer gas. Electron-based dissociation (EXD where X = C for ECD or X = I for EID) was performed in Q5, which is equipped with an axial electron source enabling interaction of trapped ion cloud with a pulsed electron beam of controlled kinetic energy. The reaction time was defined by the electron irradiation period while ions remained radially and axially confined. The electron kinetic energy was adjusted according to the selected dissociation mode, typically ∼0 eV for ECD and ∼35 eV for EID. Collisionally activated ECD (ECciD) was performed by combining ECD with supplemental _R_CID.

Modular multimodal MS^3^ workflows were performed by combining isolatiom, activation, transger and re-isolation steps. In the present experiments, MS^2^ product ions generated by _R_CID or ECD were isolated and subjected to a second activation step by ECD or _R_CID, enabling sequential MS^3^ combinations. Product ions could optionally be accumulated in parallel in downstream segments (Q8) to enhance ion statistics prior to ejection (ion enrichment workflow).

Precursor ions were accumulated for 30 ms to 1 s prior to activation, depending on precursor abundance. For electron-based dissociation experiments, irradiation times were adjusted according to precursor intensity and desired fragmentation efficiency. For _R_CID experiments, excitation amplitude and duration were optimized independently. Acquisition parameters for each MS^2^/MS^3^ spectrum (accumulation times, irradiation times, electron energy, excitation settings and isolation settings) are indicated directly in the corresponding figures and compiled in the supplementary tables.

### Data Analysis

Top-down MS^2^ and MS^3^ spectra were exported from Compass DataAnalysis (Bruker Daltonics) as simple ASCII (.xy) files containing m/z and intensity values. Spectra were recalibrated in OmniScape^TM^ (hereafter abbreviated as OSc) software version 2025b (Bruker Daltonics), when necessary, using the Calibration workflow. Recalibration was performed based on matched theoretical fragment ions of the most probable proteoform using a quadratic fit function. For each calibration, more than 10 calibration points were selected across the full *m/z* range.

Proteoform analysis was performed using the “Confirmation” workflow in OSc, a workflow for fragment identification for proteins with known sequence and user defined PTMs. For *in vitro* reactions, variable acetylation was set on all possible lysine residues of histone H4 and H3.1. Regarding endogeneous histone, acetylation, mono-, di- and trimethylation were set as variable PTMs on all possible lysine residues of histone H4. The software operates directly on profile *m/z* spectra, using the OmniWave^TM^ algorithm, calculating theoretical isotopic distributions for all possible charge states and selecting the optimal set of isotopic distributions that best explain the mass spectrum. Thus, for each candidate proteoform, theoretical fragment ions and their isotopic distributions are assigned with high confidence, even in highly congested top-down mass spectra. Fragment ion types follow the classical a, b, c and x, y, z nomenclature (55) with numbering from the N- and C-termini, respectively.

Fragment matches were accepted based on agreement between experimental and theoretical isotopic envelopes using the following criteria: mass tolerance 5 ppm; score threshold 3; a minimum intensity within mass tolerance (ei score) of 60-80%; a correlation threshold of 20–50% and a stringency of peak acceptance set to “High” (0.35).

Proteoform ranking relied primarily on the MS score, which quantifies the agreement between experimental and theoretical fragment ion isotopic distributions for a given proteoform hypothesis. It is derived from fragment-level matches satisfying mass accuracy and isotopic quality criteria. In addition, sequence-level validation is assessed using the Sequence Validation Percentage (SVP). SVP estimates the fraction of a protein sequence for which unexpected sequence variants or post-translational modifications can be excluded based on fragment constraints, while tolerating terminal and internal gaps where sequence length and composition remain mass constrained (56). SVP thus reflects experimentally constrained backbone coverage but is intrinsically insensitive to positional PTM isomerism when multiple proteoforms satisfy the same constraints. SVP parameters were set to a terminal gap size of two residues and an internal gap size of a single residue.

Reports generated by OSc for proteoform identification are provided in Supporting Information. Deconvoluted spectra shown in Figure 7A-ii were generated using UNIDEC (57).The following parameters were applied: *m/z* range 500-1400; charge state range 1-50, mass range 5000-25000 Da, mass sampling interval 0.2 Da; peak full width at half maximum 0.01 Th, and Gaussian peak shape function.

### Proteoform nomenclatures

Proteoforms are described in this work using bracket-based notation introduced by Kelleher and co-workers (58). Square brackets ‘[]’ indicate fully defined proteoforms in which all PTMs are chemically identified and site-localized. For example, [H3.1K14ac] denotes histone variant H3.1 acetylated exclusively at K14. Curly braces ‘{ }’ denote partially defined proteoforms, where the presence of at least one or more PTMs is established but their localization remains unresolved. For example, {H4Kac2} indicates a histone H4 carrying two lysine acetylation without specification of the modified lysine residues.

### Experimental Design and Statistical Rationale

This study was designed as an analytical evaluation of multimodal top-down MSⁿ workflows on the timsOmni platform for characterization of intact histone proteoforms. Recombinant histones H3.1 and H4 were subjected to independent *in vitro* acetylation reactions with GCN5, PCAF, or p300, and a liver cell fraction containing endogenous histone H4 was also analyzed. The aim was to assess how complementary ion activation methods (_R_CID, ECD, EID, and ECciD) and targeted MS³ experiments improve sequence coverage, PTM localization, and discrimination of positional isomers.

No formal statistical hypothesis testing was applied. Proteoform assignments were based on analytical criteria implemented in OSc, including fragment-isotope matching, mass accuracy, MS score, and SVP. Supplementary Tables S1-S6 summarize the precursor m/z selected and the MS score, SVP, SVP range, and sequence coverage (%) for the top-ranked candidate proteoforms identified by OSc. OSc reports are also provided in the Supporting Information for every experiment presented.

## Results and discussion

### GCN5-catalyzed acetylation of histone H3.1

Figure 2 presents the results of a ten-minute *in vitro* reaction of histone variant H3.1 performed in the presence of the histone acetyltransferase GCN5 (KAT2A). The MS¹ signal of H3.1 was dominated by a single monoacetylated species {H3.1Kac} (13,315 Da), together with its oxidized form (13,331 Da) (Figure 2A). MS/MS spectra were acquired for 19+ and 20+ precursor ions that using _R_CID, ECD, EID, and ECciD (Figure 2B).

**Figure 1.**
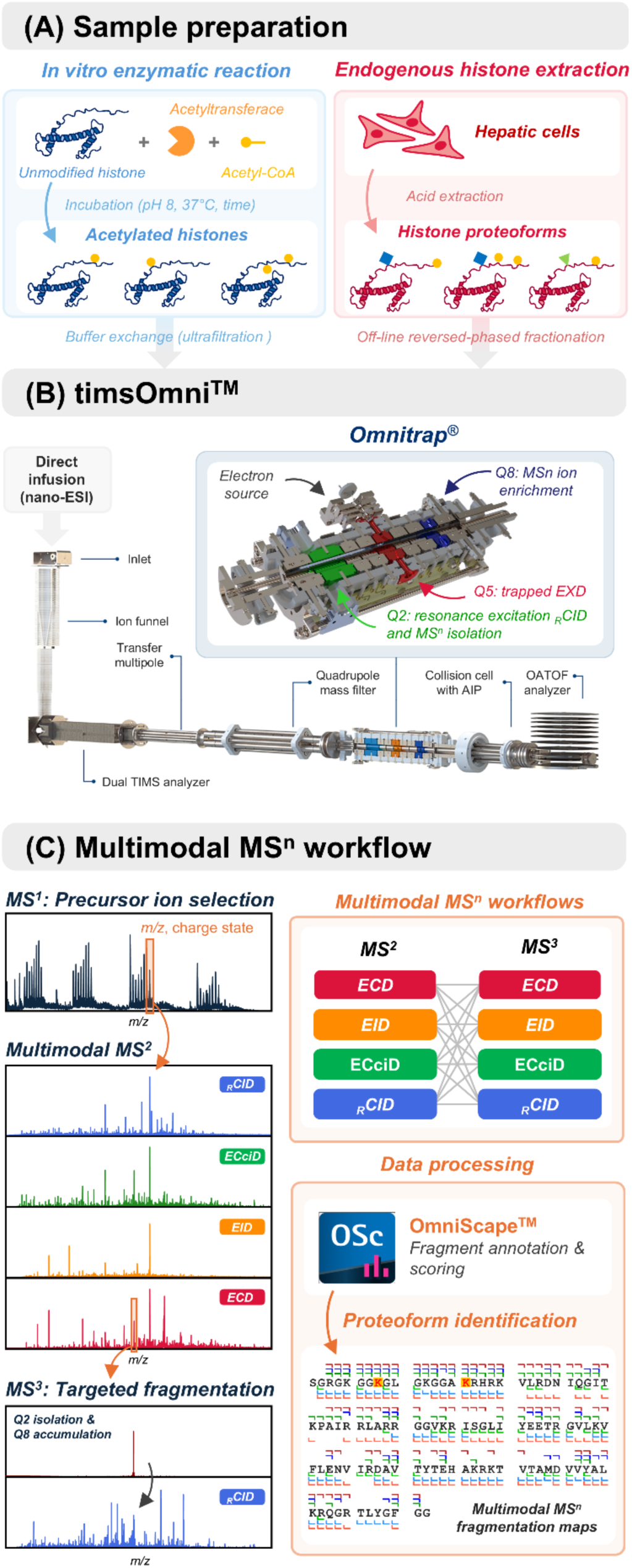
Overview of sample preparation and multimodal MSⁿ workflow for top-down analysis of histone proteoforms. (A) Histone H4 and H3.1 proteoforms were generated by *in vitro* enzymatic acetylation reactions using p300, PCAF, GCN5 while endogeneous H4 was isolated from hepatic cells by acid extraction followed by reversed-phase micropurification. (B) Schematic of the timsOmni prototype integrating an Omnitrap platform, which enables multiple ion activation modes. (C) Multimodal MSⁿ workflow. Precursor ions of specific m/z and charge state are isolated and subjected to complementary MS^2^ fragmentation (RCID, ECD, EID, and ECciD). Targeted MS^3^ experiments can be performed on selected product ions to refine PTM localization. Fragment ions are annotated and scored using OmniScape, yielding multimodal fragmentation maps for proteoform identification.

**Figure 2.**
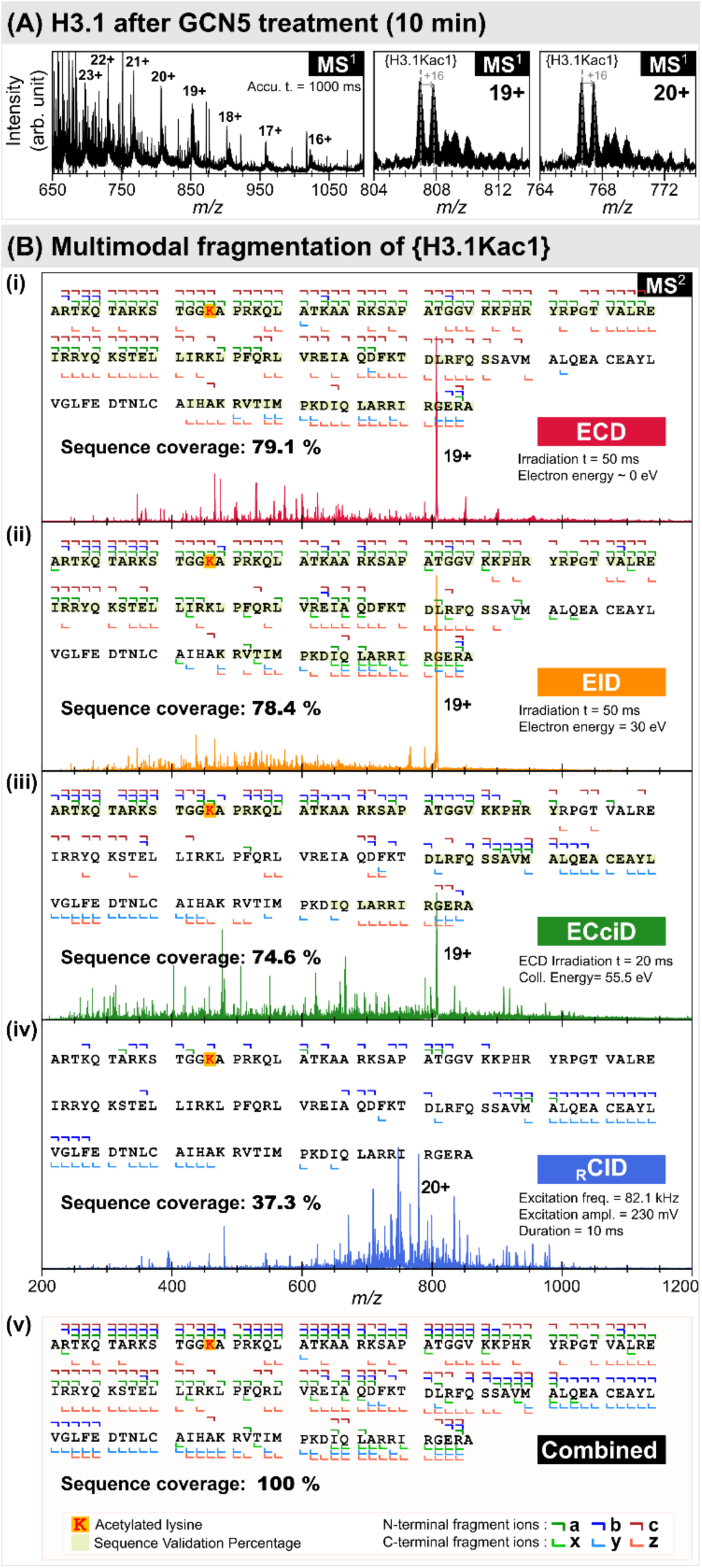
Multimodal fragmentation analysis of monoacetylated H3.1 produced by GCN5. (A) MS^1^ spectrum of histone H3.1 after a 10 min i*n vitro* reaction with GCN5. (B) Multimodal MS^2^ fragmentation of the identified proteoform [H3.1K14ac]. (i–iv) Representative spectra obtained by ECD, EID, ECciD (19^+^ precursor) and RCID (20^+^ precursor), with corresponding fragment assignments along the H3.1 sequence. (v) Combined fragmentation map summarizing fragment ions identified across all datasets.

Across fragmentation modes, the acetylation was consistently localized to K14 (Table S1), and electron-based methods (ECD, EID and ECciD) provided individual sequence coverages ranging from 66.4 to 79.1% (Figure 2B(i-iii)). The identification of [H3.1K14ac] was based on the multiple diagnostic a-, b- and c-type fragments ions carrying a +42 Da mass shift in the 14-17 residues region. In addition, [H3.1K18ac] and [H3.1K9ac] were excluded from further consideration, as they were supported only by sparse, low intensity fragment ions that were not reproduced across all conditions. _R_CID alone provided lower sequence coverages, with 41.8 % and 37.3 % for the charge states 19+ and 20+, respectively. Nevertheless, both datasets remained consistent with acetylation at K14, which gave the highest MS score, although SVP was limited to the residues 1-6 and 132-135 (Figure 2B(iv)). The lower sequence coverage likely resulted from the preferential cleavage of labile bonds during resonance activation. When all the different fragmentation data were combined, the sequence coverage reaches 100% (Figure 2B(v)), with the best continuous SVP across residues 1-86 and 112-135 obtained for ECD, covering all possible lysine residues.

To confirm K14 localization, an MS³ experiment was performed in which ECD of {H3.1Kac1}^19+^ produced a charge-reduced radical intermediate (18+•), followed by _R_CID (Figure S1A). This increased MS^3^ sequence coverage to 56%, representing a ∼35–50% gain relative to direct MS²(_R_CID) (Figure S1B(i)). The SVP score also increased from 6.7% to 17.8%. Fragments spanning residues 1-17 localized the acetylation to K14 while excluding modification at K9 and K18, confirming that [H3.1K14ac] was the predominant proteoform. Overall, application of multimodal MS^n^ workflows confirmed that GCN5 predominantly acetylated H3.1K14, while no other major acetylation sites were detected, in agreement with the previously reported preference of GCN5 for H3K14(59).

### PCAF-catalyzed acetylation of histones H3.1 and H4

PCAF (KAT2B) displays a well-established preference for histone H3.1, whereas histone H4 is a comparatively weaker substrate (60–62). To evaluate enzyme specificity at the intact proteoform level, separate 30 minutes *in vitro* acetylation reactions were performed with recombinant H3.1 and H4.

The MS^1^ spectrum of H3.1 after PCAF treatment is presented in Figure 3A(i). A zoomed view of the region of the charge state 19+ (Figure 3A(ii)) showed only acetylated species, with no detectable unmodified H3.1 ([H3.1]). The monoacetylated precursor {H3.1Kac1}^19+^ was selected for ECD, EID and _R_CID characterization. Across all activation modes, [H3.1K14ac] was identified as the top-ranked proteoform, followed by [H3.1K9ac] and [H3.1K18ac] (Table S2). In ECD spectra, [H3.1K14ac] gave the highest MS score (28.7) and the largest SVP (67.4%), with N-terminal sequence validation extending from residues 1-64. By contrast, although [H3.1K18ac] ranked second (MS score 27.1), its SVP dropped to 29.6%, restricted to residues 1-13 in the N-terminal region. Moreover, no unmodified N-terminal fragments were detected in region 14-17, supporting localization of the acetylation at or before K14, thus excluding K18. No fragment ions provided evidence for acetylation at K4, while support for [H3.1K9ac], ranked third (MS score 26.7), was supported by SVP validation up to residue 10 and with two low-intensity acetylated c_9_^2+^ and c_10_^2^+ fragments.

**Figure 3.**
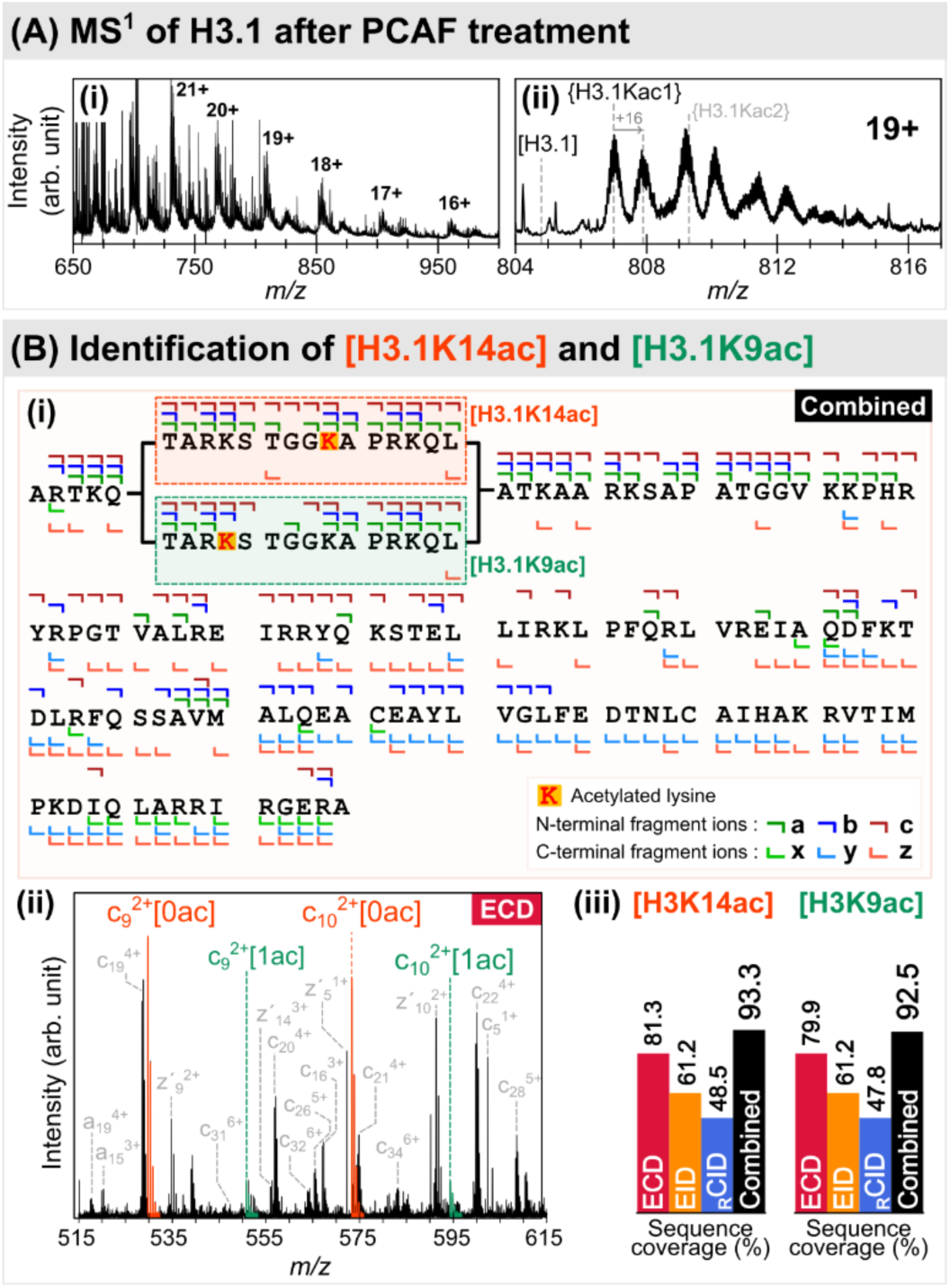
Identification of monoacetylation sites in histone H3.1 after PCAF treatment using multimodal MS^2^ analysis. (A) (i) MS^1^ spectrum of H3.1 after a 30 min *in vitro* acetylation reaction with PCAF. (ii) Zoomed view of the charge-state 19⁺ region showing the monoacetylated population. (B) Characterization of the monoacetylated species {H3.1Kac1}. (i) Combined fragmentation map summarizing fragment ions identified using ECD, EID, and RCID. (ii) Zoomed ECD spectrum showing diagnostic unmodified ([0ac]) and monoacetylated ([1ac]) of c_9_^2+^ and c_10_^2+^ ions, supporting the dominant [H3.1K14ac] and minor [H3.1K9ac] proteoforms. (iii) Sequence coverage obtained for each individual fragmentation method (colored bars) and for the combined dataset (black bars) for both identified proteoforms.

Based on the relative intensities of the acetylated and unmodified forms of c_9_^2+^ and c_10_^2+^ fragment ions, and assuming comparable ionization, fragmentation and detection efficiencies, the relative abundance of [H3.1K14ac] versus [H3.1K9ac] was estimated to be approximately 7.1 and 9.5 (Figure 3B(ii)). This suggests that [H3.1K14ac] was the major monoacetylated proteoform, accounting for roughly 90% of the population and [H3.1K9ac] represented a minor fraction of about 10%. These values should be regarded as rough estimates intended to reflect a relative order of magnitude rather than absolute quantification.

Among the different MS² techniques tested, ECD provided the highest sequence coverage (81.3%) for [H3.1K14ac], while EID and _R_CID offered complementary coverage (61.2% and 48.5%, respectively) (Figure 2B(iii)). Combined all fragmentation modes increase the total sequence coverage to 93.3% for the [H3.1K14ac] and 92.5% for [H3.1K9ac]. The MS^1^ spectrum also suggests the presence of a diacetylated H3.1 species (Figure 3A(ii)), but the spectral quality in this region did not allow acquisition of MS/MS data sufficient for confident localization of a second acetylation site.

The activity of PCAF toward histone H4 was examined under identical reaction conditions (Figure 4). The total MS¹ spectrum showed a dominant monoacetylated species, {H4Kac1}, and a lower-abundance doubly acetylated species, {H4Kac2}, together with residual unmodified H4 (Figure 4A right panel). No clear signals were detected at the expected positions for {H4Kac3} or {H4Kac4} (light grey annotation). Instead, several low-intensity peaks appear consistent with adducted and oxidized forms of the protein.

**Figure 4.**
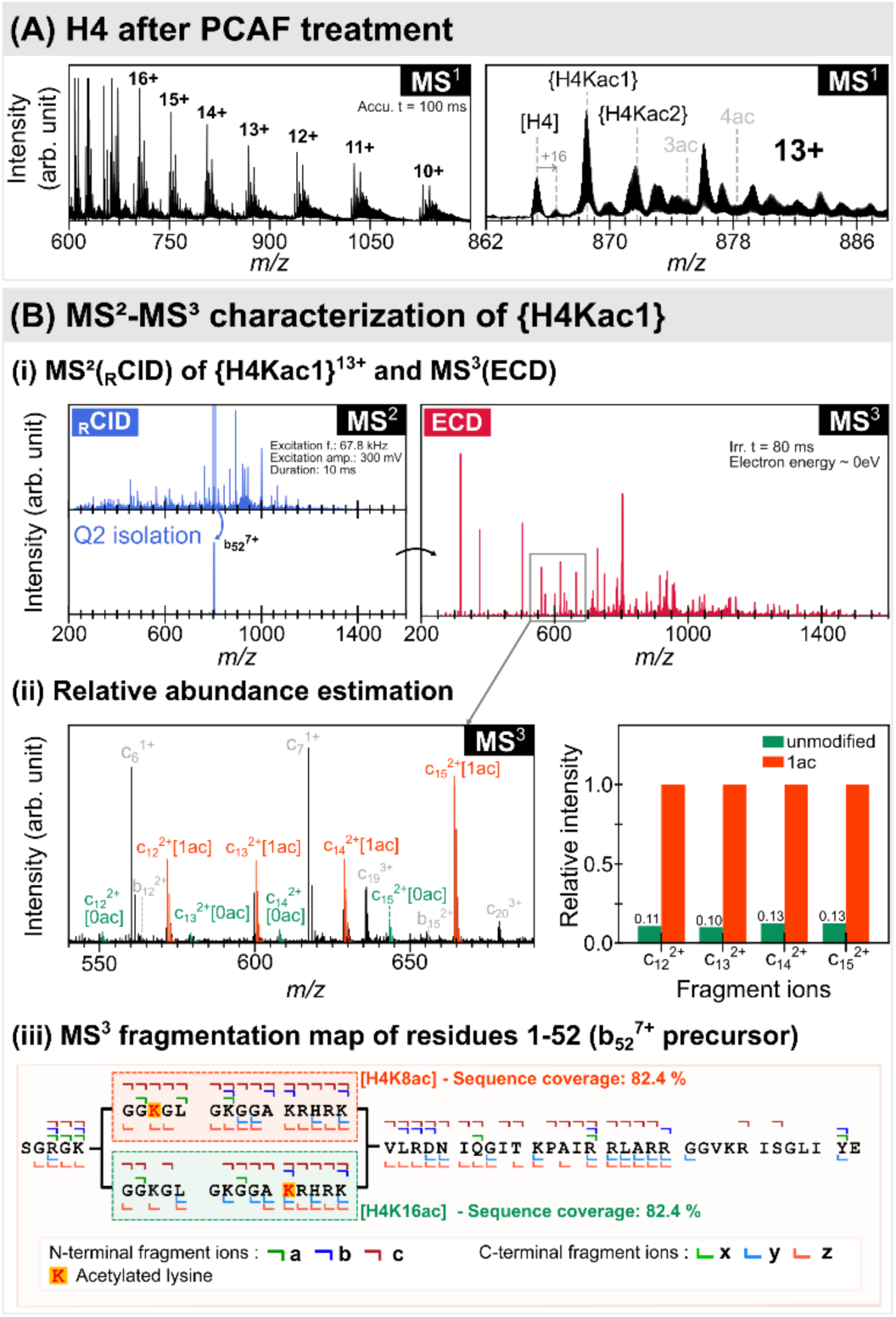
Localization of monoacetylation sites in histone H4 after PCAF treatment using targeted MS3 analysis. (A) MS^1^ spectrum of H4 after *in vitro* acetylation by PCAF for 30 min, showing the charge-state distribution and a zoomed view of the 13^⁺^ precursor region containing a dominant {H4Kac1} population. (B) (i) targeted MS2(RCID)→MS3(ECD) workflow applied to {H4Kac1}13^+^. The N-terminal b52^7+^ fragment (residue 1-52) bearing one acetylation, was isolated in Q2 and accumulated (left spectra) before MS^3^ analysis by ECD (right spectrum). (ii) Left: zoomed view of diagnostic unmodified ([0ac]) and monoacetylated ([1ac]) c-type fragment ions (c12-15^2+^). Right: histogram of the corresponding relative fragment intensities. (iii) Fragmentation map and PTM localization within the H4 N-terminal tail for the main ([H4K8ac]) and secondary ([H4K16ac]) proteoforms.

The monoacetylated precursor {H4Kac1}^13+^ was first analyzed by MS^2^ using ECD, EID, ECciD and _R_CID. Across all activation modes, [H4K8ac] was consistently ranked as the top candidate proteoform, followed by [H4K12ac], while [H4K16ac] and [H4K5ac] ranked third and fourth depending on the fragmentation method (Table S3). However, N-terminal c-type ions spanning residues 5-8 were detected only in their unmodified forms, thus excluding [H4K5ac] despite its comparable MS score.

To refine localization within the N-terminal tail, the abundant _R_CID fragment b ^7+^ was mass-selected and subjected to MS^3^(ECD) analysis (Figure 4B). In this experiment, [H4K8ac] remained the top-ranked proteoform (MS score 28.1, SVP 82.4%, supported by SVP across N-terminal residues 1-40) with [H4K12ac] and [H4K16ac] ranked second and third, respectively (MS score 26.1 and 25.0). The c_8_-c_11_ ions were only detected unmodified therefore did not providing evidence for [H4K12ac]. By contrast, low intensity diagnostic fragments (acetylated and unmodified c_12-15_^2+^ and y_y39_^3+^), supported the presence of [H4K16ac]. Comparison of these shared fragment-ion intensities suggested a [H4K8ac]:[H4K16ac] abundance ratio of about 8.6 ± 1.0 (Figure 4B(ii)).

Taken together, the indicated that {H4Kac1} predominantly corresponded to [H4K8ac] under the present *in vitro* conditions, with a minor contribution from [H4K16ac]. Combined MS² and targeted MS³ data yielded 99% sequence coverage for the dominant assignment.

A lower-abundance doubly acetylated population, {H4Kac2} was also detected (Figure 4A), although partially overlapped with isobaric/isomeric background signals. MS^2^ (ECD and _R_CID) produced several closely ranked candidate proteoforms (Table S4). Targeted MS³(ECD) experiments (Figure S2B(i)) was performed on the doubly acetylated _R_CID b ^7+^ fragment consistently supported acetylation at K8 within the {H4Kac2} population. Site-specific fragments further supported [H4K8acK16ac] and [H4K8acK44ac], (Figure S2B(ii-iii)), whereas intermediate lysines (K20 and K31) remained plausible but cannot be uniquely assigned. A detailed description of the MS³ analysis is provided in the Supporting Information.

The multimodal MS^n^ workflow demonstrates that PCAF acetylates both histones H3.1 and H4 *in vitro*, with distinct levels of site-specificity. On monoacetylated H3.1, PCAF showed a predominant preference for K14, with a minor contributions from K9, whereas on monoacetylated H4 was assigned mainly to [H4K8ac] with minor [H4K16ac]. These results are consistent with previous studies reporting H3K14 as the primary PCAF target with minor acetylation at H3K9, and H4K8 as preferred site on H4 with minor acetylation at H4K16 (62–64). In contrast, the doubly acetylated H4 population showed greater positional heterogeneity but K8 remains an anchor site, while the second acetylation was distributed across multiple possible lysines residues with K16 and K44 directly supported.

### p300-catalyzed multiple acetylation of histone H4

The histone lysine acetyltransferase p300 (KAT3B) is a transcriptional co-activator with broad substrate specificity, acetylating all four core histones (H2A, H2B, H3, and H4), as well as numerous non-histone proteins *in vivo* (65). In contrast to the more site-selective acetyltransferases GCN5 and PCAF, p300 can modify multiple lysine residues within the histone tails. On histone H4, reported sites include the N-terminal tail K5, K8, K12, and K16 with a preference for K5 and K8, together with additional lysine residues have also been reported at lower abundance (62, 66, 67). Consistent with this broad specificity, incubation of H4 with p300 for 1h produced a wide distribution of acetylated species (Figure 5A), ranging from unmodified H4 (11,236 Da) to proteoforms carrying up to seven acetylations. Subsequent analysis focused on {H4Kac6} and {H4Kac7}, as these highly acetylated species provide an informative model for localization of multiple co-occurring acetylation sites on intact histones.

**Figure 5.**
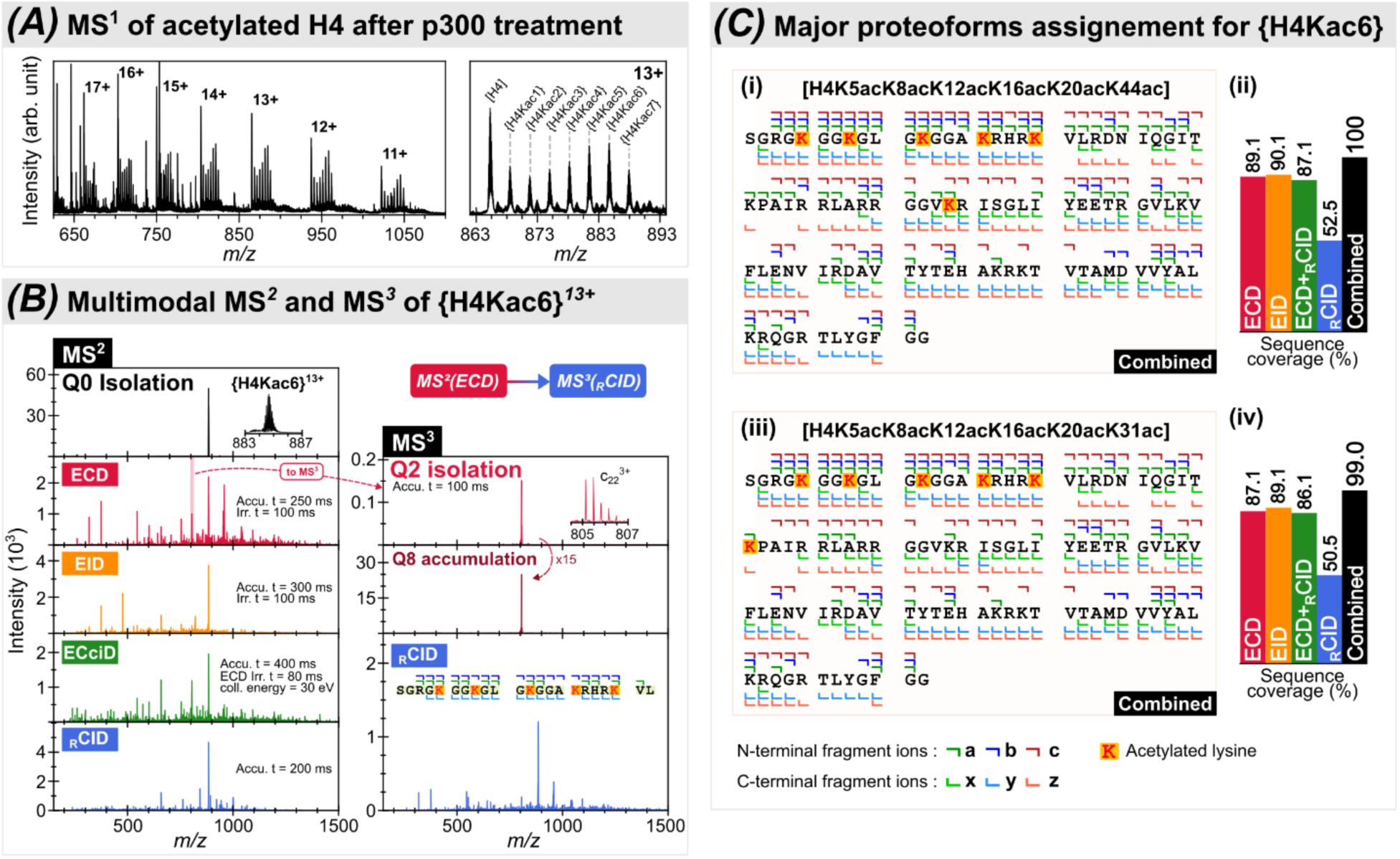
Proteoform characterization of hexa-acetylated histone H4 ({H4Kac6}) generated by p300 using a multimodal MSⁿ workflow. (A) MS1 spectrum of H4 after 1h *in vitro* acetylation by p300, showing the charge-state distribution (11+-17+) and zoomed view of the 13^⁺^ region. (B) Multimodal MS² and targeted MS³ workflow applied to the {H4Kac6}^13+^ precursor. Left panels: MS^2^ spectra acquired using ECD, EID, ECciD, and RCID. Right panels: targeted MS^3^ experiment based on ECD followed by RCID after isolation of the c22^3+^ fragment ion in Q2 and accumulation in Q8. (C) Proteoform assignment for {H4Kac6}. (i, iii) Combined fragmentation maps showing identified fragment ions (a, b, c and x, y, z series) and localized acetylated lysine residues for the two main proteoforms. (ii, iv) Sequence coverage obtained for each fragmentation method (ECD, EID, ECciD, RCID) and for the combined dataset.

Figure 5B summarizes the multimodal MS^n^ workflow applied to for {H4Kac6}^13+^. ECD of the intact precursor generated a relatively abundant c_22_^3+^ fragment corresponding to the H4 N-terminal tail, which was isolated in Q2 for 100 ms and accumulated in Q8, resulting in a ∼15-fold signal enhancement. Subsequent _R_CID on this fragment yielded 95.2% sequence coverage of the tail region and showed that major {H4Kac6} population carried five N-terminal acetylation sites at K5, K8, K12, K16 and K20, thereby constraining the remaining acetylation to the downstream sequence.

Multimodal MS^2^ analysis provide complementary information to localize the sixth site. [H4K5acK8acK12acK16acK20acK44ac] was the top-scoring proteoform (Table S5), while [H4K5acK8acK12acK16acK20acK31ac] was ranked second, in the EXD datasets. ECD, EID and ECciD provided sequence coverage of 89.1%, 90.1%, and 87.1% for [H4K5acK8acK12acK16acK20acK44ac], with an MS score of 54.7, 35.3 and 40.8, and N-terminal sequence validation extending to residue 41, 46 and 30, respectively. In the ECD spectrum, a continuous c-ion series spanning residues 33-41 were observed with either five or six acetylations, consistent with coexisting [H4K5acK8acK12acK16acK20acK44ac] and [H4K5acK8acK12acK16acK20acK31ac]. _R_CID data supports presence of the [H4K5acK8acK12acK16acK20acK44ac] yielding 52.5% sequence coverage through diagnostic b- and y-type fragments (b_24_^4+^, y_63_^7+^, y_49_^5+^ and y_47_^4+^). In addition, two low-intensity c_22_^3+^ and c_23_^3+^ ions carrying four and five acetylations suggested a third minor configuration, [H4K5acK8acK12acK16acK31acK44ac], although this proteoform is not supported by ECciD, suggesting that its abundance is near detection limit. No consistent fragment evidence supports acetylation beyond K44.

The relative abundances of the two principal proteoform, [H4K5acK8acK12acK16acK20acK44ac] and [H4K5acK8acK12acK16acK20acK31ac], were estimated from ECD data, which provide the largest number of diagnostic fragments, using shared c_34_ and c_36_ to c_41_ ions observed with five or six acetylations together with complementary z_59_ to z_67_ bearing one and no acetylation at identical charge states. The average abundance ratio between these two proteoforms is 1.8 ± 0.5, indicating that {H4Kac6} was dominated by [H4K5acK8acK12acK16acK20acK44ac] (∼70%), followed by [H4K5acK8acK12acK16acK20acK31ac] (∼25%). The possible configuration [H4K5acK8acK12acK16acK31acK44ac] accounted for only a minor fraction (∼5%). ECciD data yields abundance ratios of 1.9 ± 0.6 between the two main proteoforms, estimated based on c_34_^5+^, c_36_^5+^-c_41_^5+^, z’_59_^6+^-z’_64_^6+^ and z_66_^6+^, which consistent with ECD-derived estimates. Integration of the multimodal MS^n^ workflow yielded overall in sequence coverages of 100% and 99% for [H4K5acK8acK12acK16acK20acK44ac] and [H4K5acK8acK12acK16acK20acK31ac], respectively (Figure 5C).

The most heavily modified species, {H4Kac7}^13+^, was characterized by ECD and _R_CID (Figure S3). Both methods identified [H4K5acK8acK12acK16acK20acK31acK44ac] as the top-scoring proteoform. and restricted the hepta-acetylated population to two positional isomers differing by acetylation at either K44 or K59, with the K44ac configuration dominating (∼75-80%). Combined ECD and _R_CID sequence coverages lead to 95.0% and 93.1% sequence coverage of for [H4K5acK8acK12acK16acK20acK31acK44ac] and [H4K5acK8acK12acK16acK20acK31acK59ac], respectively. Detailed analysis is provided in the Supporting Information.

### Evolution of sequence coverage during data acquisition

To evaluate how backbone information accumulates with increasing data acquisition, sequence coverage was analyzed as a function of the number of accumulated scans, N (Figure 6). For each value of N, at least three subsets were evaluated when possible, and sequence coverage is reported as mean ± standard deviation. The analysis was restricted to datasets dominated by a single proteoform to minimize bias from coexisting positonal isomers.

**Figure 6.**
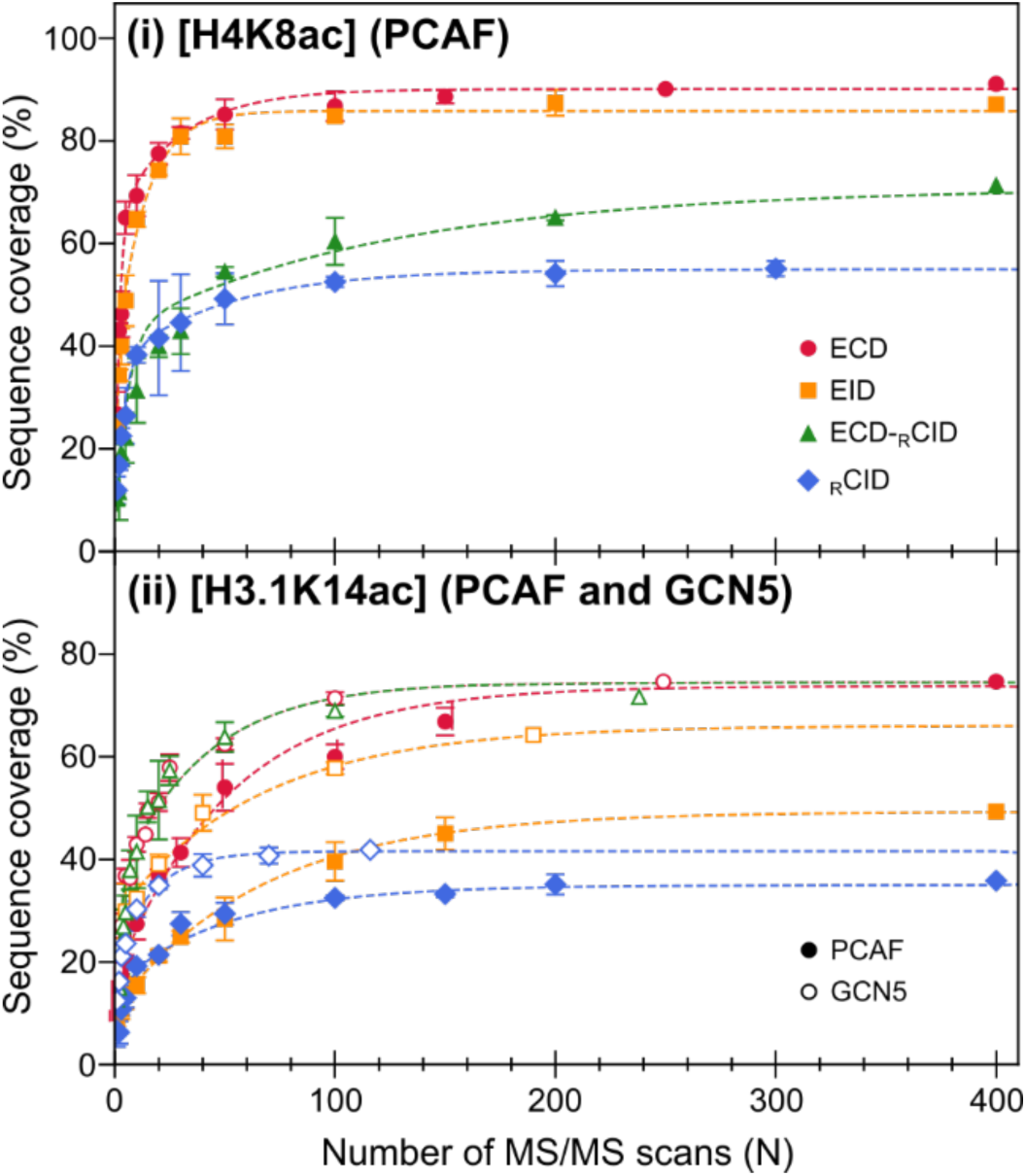
Sequence coverage as a function of the number of accumulated MS/MS scans (N) for (i) [H4K8ac] generated by PCAF and (ii) [H3.1K14ac] generated by PCAF and GCN5. Experimental values are shown as mean ± standard deviation (n = 3). Fragmentation methods are indicated by color and symbol: ECD, red circles; EID, orange squares; ECciD, green triangles; and RCID, blue diamonds. In panel (ii), filled symbols correspond to PCAF-treated samples and open symbols correspond to GCN5-treated samples. Dotted lines represent fits of the data using a bi-exponential model.

Sequence coverage was expressed as a function of the number of accumulated MS/MS scans rather than acquisition time because scan duration depends on experimental settings, in particular precursor accumulation, and varied from 30 to 1,000 ms per scan. Under these conditions, N provides a more comparable metric for direct comparison across different experiments.

For all fragmentation methods, sequence coverage increased rapidly at low scan numbers and then more gradually as additional scans were accumulated. This behavior reflects the early detection of the most abundant fragment ions arising from the most favored backbone cleavages, while less probable cleavages required accumulating higher number of scans to be detected. The dependence of sequence coverage on N was described empirically using a bi-exponential saturation function (dotted lines in Figure 6):

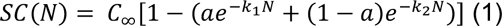

Where *SC*(*N*) is the sequence coverage obtained from N spectra, *C*_∞_ is the asymptotic plateau, k1 and k2 describe fast a slow accumulation regime, and *a* is the fractional contribution of the fast component. Parameters were estimated by nonlinear least-squares minimization.

The bi-exponential model provided excellent agreement with the data (R² = 0.9980–0.9999, mean 0.9995), whereas a single-exponential model resulted in lower goodness-of-fit (R² = 0.775–0.968, mean 0.918), and systematically overestimated sequence coverage at low scan numbers while underestimating the plateau level (Figure S4). Based on the bi-exponential fits, electron-based fragmentation reached high sequence coverage after relatively few scans, typically 2-4 ECD and EID scans for H4 to reach 50% sequence coverage, 5-20 scans for H3.1 depending on the activation method and precursor. _R_CID showed a similar behavior but reached lower plateau values. The faster convergence observed for H4 likely reflects a combination of higher precursor signal intensity and the shorter H4 sequence, which reduced the number of backbone cleavages required to reach a given sequence coverage.

### Characterization of endogenous human histone H4 proteoforms

Figure 7A shows the MS¹ (i) and deconvoluted (ii) spectra of a histone fraction extracted from human liver cells after off-line reversed-phase fractionation. Histone H4 was a major component, together with signals in the mass range expected for H2B (13,700-13,900 Da) and H2A (13,950-14,100 Da). The deconvoluted spectrum reveals two major H4 species at 11,306 and 11,348 Da, corresponding to +70 Da and +112 Da relative to unmodified H4 (11,236 Da). Minor neighboring peaks differing by +14 Da were also observed, consistent with methylation. Precursor ions at *m/z* 707 and 710 (16+) were mass-selected for multimodal MS² analysis by ECD, EID, and _R_CID.

**Figure 7.**
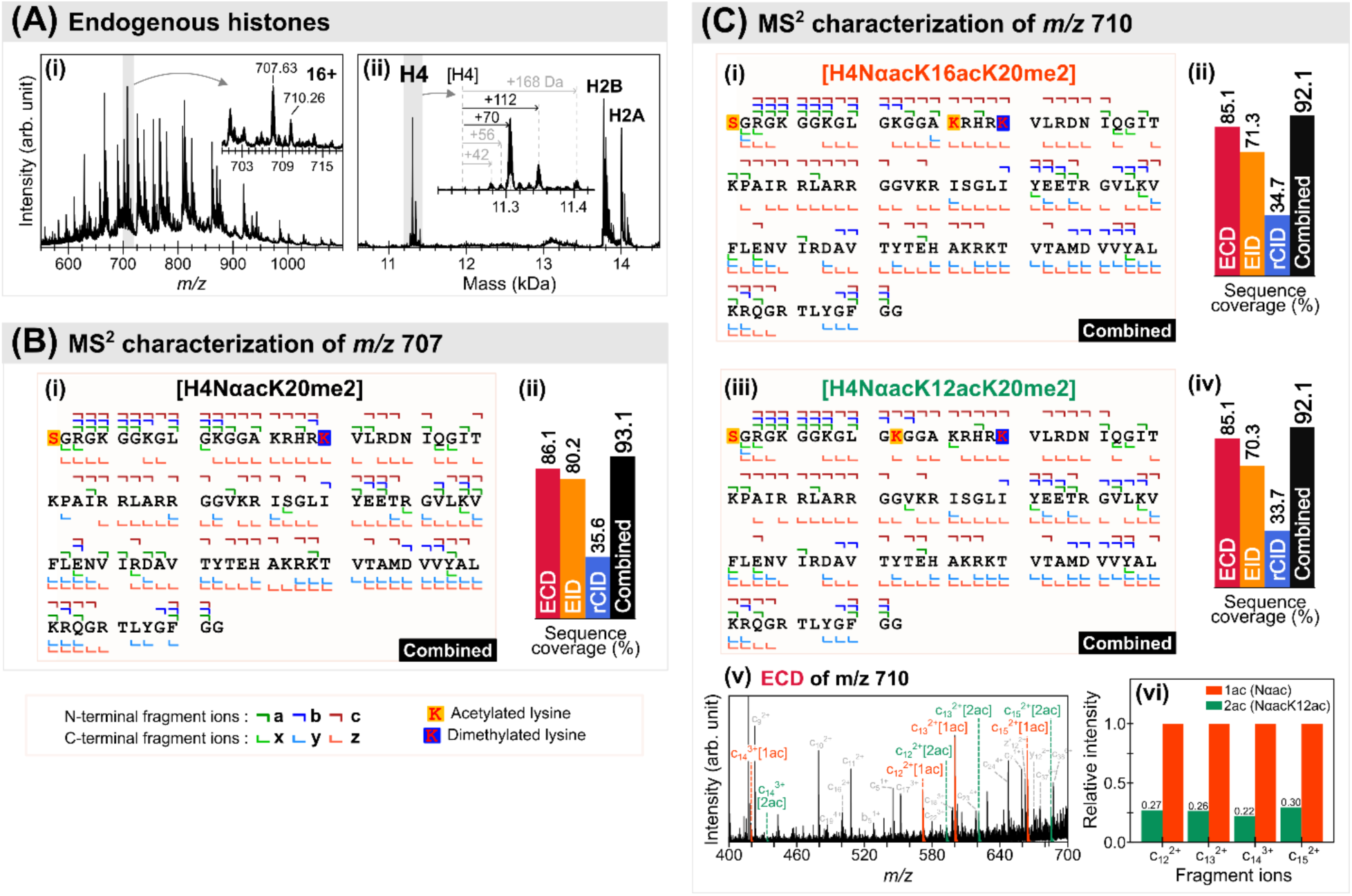
Characterization of endogenous histone H4 proteoforms using multimodal MS^2^ analysis. (A) MS^¹^ spectrum (i) and deconvoluted mass spectrum (ii) of an offline histone fraction extracted from liver cells. (B) Multimodal MS^2^ analysis of the precursor ion at *m/z* 707 (16^+^). (i) Combined fragmentation map showing identified a-, b-, c-, x-, y- and z-type fragment ions and annotated PTMs. (ii) Sequence coverage obtained for each fragmentation method (ECD, EID and RCID) and for the combined dataset. (C) Multimodal MS^2^ analysis of the precursor ion at m/z 710 (16^+^). (i, iii) Combined fragmentation maps corresponding to two candidate proteoforms, showing identified fragment ions and PTM annotations. (ii, iv) Sequence coverage obtained for each fragmentation method (ECD, EID and RCID) and for the combined dataset. (v) Zoomed view of the ECD spectrum highlighting selected diagnostic fragment ions. (vi) Relative intensities of the corresponding diagnostic fragment ions.

For the *m/z* 707 precursor, ECD and EID consistently supported a doubly modified proteoform carrying one +42 Da modification and one +28 Da modification. In both datasets, the two top-ranked proteoforms consistently identify by OSc were [H4NαacK20me2] and [H4K5acK20me2] (Table 6). These assignments differ only in the position of the acetylation, located either at the protein N terminus or at K5, while both place a dimethylation at K20 (K20me2). N-terminal fragment ions up to residue 4 were consistently detected only in their acetylated form, supporting N-terminal acetylation (Nαac). The alternative K5ac assignment lacked consistent corroboration across the different MS/MS experiments and was supported only by aa single low-intensity ion (a_3_^1+^) in EID dataset. No fragment ions supported lysine modifications at K8, K12, or K16. For the additional +28 Da shift fragment ions do not support the presence of two monomethylated lysine residues. Fragment ions spanning the K20-K31 region carried both modifications, supporting dimethylation at K20 rather than two monomethylation on two different lysine residues. _R_CID also ranked [H4NαacK20me2] as the top proteoform, although with lower sequence coverage (33.7%). Taken together, [H4NαacK20me2] remains as the only proteoform identified for the *m/z* 707 precursor, with an overall sequence coverage of 93.1% (Figure 6B(ii)).

The *m/z* 710 precursor, Osc and analysis of fragment ion indicated a modification pattern consistent with the *m/z* 707 species carrying one additional acetylation. ECD fragment-ion analysis localized one the second at K16, thus supporting the presence of [H4NαacK16acK20me2] with a sequence coverage of 85.1 % and 71.3 % using ECD and EID, respectively (Figure 6C(ii)). In addition, ECD revealed lower intensity diagnostic c-type fragment ions between K12 and K16 carrying two acetyl groups (c_12_-c_15_), consistent with a minor [H4NαacK12acK20me2] positional isomer (Figure 6C(v)). EID does not provide clear evidence for this [H4NαacK12acK20me2] isomer, likely reflecting its lower sequence coverage and reduced fragment ion signal relative to ECD under the selected conditions. No additional proteoform candidates are supported by the observed fragment ion series. Based on the relative intensities of c_12_-c_15_ ions, [H4NαacK16acK20me2] was estimated to be ∼3.8-fold more abundant than [H4NαacK12acK20me2] (Figure 6C(vi)).

Because a +42 Da mass shift is compatible with both acetylation and trimethylation events, an important ambiguity could arise from these near-isobaric PTMs. This ossibility was considered for the endogenous H4 proteoforms. Indeed, these two modifications differ in mass by only 36.4 mDa. corresponding to a relative mass difference of ∼3.2 ppm for intact H4, which can be insufficient to exclude both possibilities based on precursor mass alone. However, at the fragment level, the same absolute mass difference corresponds to a larger relative difference, reaching ∼7.4 ppm for 5 kDa fragments and increasing further for smaller fragments. This exceeds both the 5 ppm mass tolerance applied for analysis and the experimental fragment mass error. Indeed, as an example for the ECD datasets of the *m/z* 707 and 710 precursor, fragment mass errors were centered around zero, with means errors (µ) of 0.7 and 0.1 ppm and standard deviations (σ) of 1.6 and 1.3 ppm, respectively (Figure S7). These values support confident distinction between acetylation and trimethylation at the fragment level and explain why OSc favored acetylated rather trimethylated candidates for the endogenous H4 proteoforms identified.

Overall, the endogeneous H4 proteoforms identified here, [H4NαacK20me2], [H4NαacK16acK20me2] and [H4NαacK12acK20me2], are consistent with the known H4 PTM landscape, dominated by N-terminal acetylation, acetylation within the K5-K16 region, and prevalent dimethylation at K20 (68–72).

## Conclusion

The timsOmni platform enables multimodal MSⁿ workflows for detailed top-down characterization of intact proteoforms. Integration of the Omnitrap enabled flexible ion trapping, isolation, and complementary ion activation by _R_CID, ECD, EID, and hybrid workflows, producing dense and complementary fragment ion series for unambiguous proteoform assignment and PTM localization. In monoacetylated H3.1, the dominant proteoform generated by GCN5 and PCAF was consistently identified as [H3.1K14ac], in agreement with known enzyme specificity. For H4, multimodal MS² and targeted MS³ experiments identified two monoacetylated proteoforms [H4K8ac] and [H4K16ac] after PCAF treatment, and multiple co-occuring acetylations and resolved site ambiguity in highly modifies species generated by p300 {H4Kac6} and {H4Kac7}. The p300 reaction generates a broad distribution of highly acetylated proteoforms yielding a more extreme combinatorial scenario than typically encountered *in vivo*, providing a particularly demanding analytical case evaluating top-down proteoform analysis. Across these experiments, complementary fragmentation approaches yielded complete or near-complete backbone sequence coverage and enabled discrimination of positional acetylation isomers in intact proteins. Targeted MS³ workflows (ECD→_R_CID, _R_CID→ECD, and charge reduction→_R_CID) were employed to resolve residual site ambiguities and to enhance local diagnostic coverage, particularly in highly modified regions. Analysis of endogenous H4 further showed that the workflow remains effective in complex biological samples and supports confident assignment of co-occurring acetylation and methylation. Together, these results highlight the capability of the timsOmni platform to support rapid multimodal MSⁿ analysis of intact proteins and residue-level mapping of combinatorial PTMs. Future developments integrating LC separation and automated data analysis will extend these workflows toward large-scale top-down proteomics and systematic characterization of proteoform complexity.

## Supporting information

Supplemental information/results

## Abbreviations

AcCoA: (acetyl-coenzyme A)
ECD: (electron-capture dissociation)
ECciD: (electron-capture collision-induced dissociation)
EID: (electron-induced dissociation)
FA: (formic acid)
HAT: (histone acetyltransferase)
IMS: (ion mobility spectrometry)
Kac: (lysine acetylation)
LC: (liquid chromatography)
MS: (mass spectrometry)
nESI: (nano-electrospray ionization)
PTM: (post-translational modification)
_R_CID: (resonance collision-induced dissociation)
SVP: (Sequence Validation Percentage)
TFA: (trifluoroacetic acid)
TIMS: (trapped ion mobility spectrometry)
TOF: (time-of-flight)

## Acknowledgment

Mass spectrometry and proteomics research at SDU in Odense, DK is supported by a generous grant to ONJ from the Novo Nordisk Foundation to establish the INTEGRA research infrastructure (grant no. NNF20OC0061575) and a generous grant from the Danish Agency of Higher Education and Science to establish PLATO: Danish National Mass Spectrometry Platform for Proteomics and Biomolecular Imaging (grant no. 5229-00012B, www.sdu.dk/PLATO). JM. acknowledges support from the Novo Nordisk Foundation (NERD grant no. NNF23OC0083407).

## Supplemental material

The supporting information document contains Supplemental Figures S1-S7 and Supplemental Tables S1-S6, including MS³ workflows, additional fragmentation maps, sequence-coverage analyses, and OmniScape ranking tables for the proteoform assignments discussed in the main text. OmniScape PDF reports are provided separately as Supplemental Data S1, containing the reports for *in vitro* acetylated recombinant histone H3.1 and H4 experiments, and reports for endogenous histone H4 analyses.

## Data availability

The data supporting this study comprises Bruker raw mass spectrometry files, exported ascii profile spectra, recalibrated spectra, and associated OSc analysis files. These materials are available from the corresponding author upon reasonable request.

