## Supplemental information/results for "Top-down Sequencing of Intact Proteoforms using the timsOmni mass spectrometer: Accurate Determination of Co-occurring Histone Modifications"

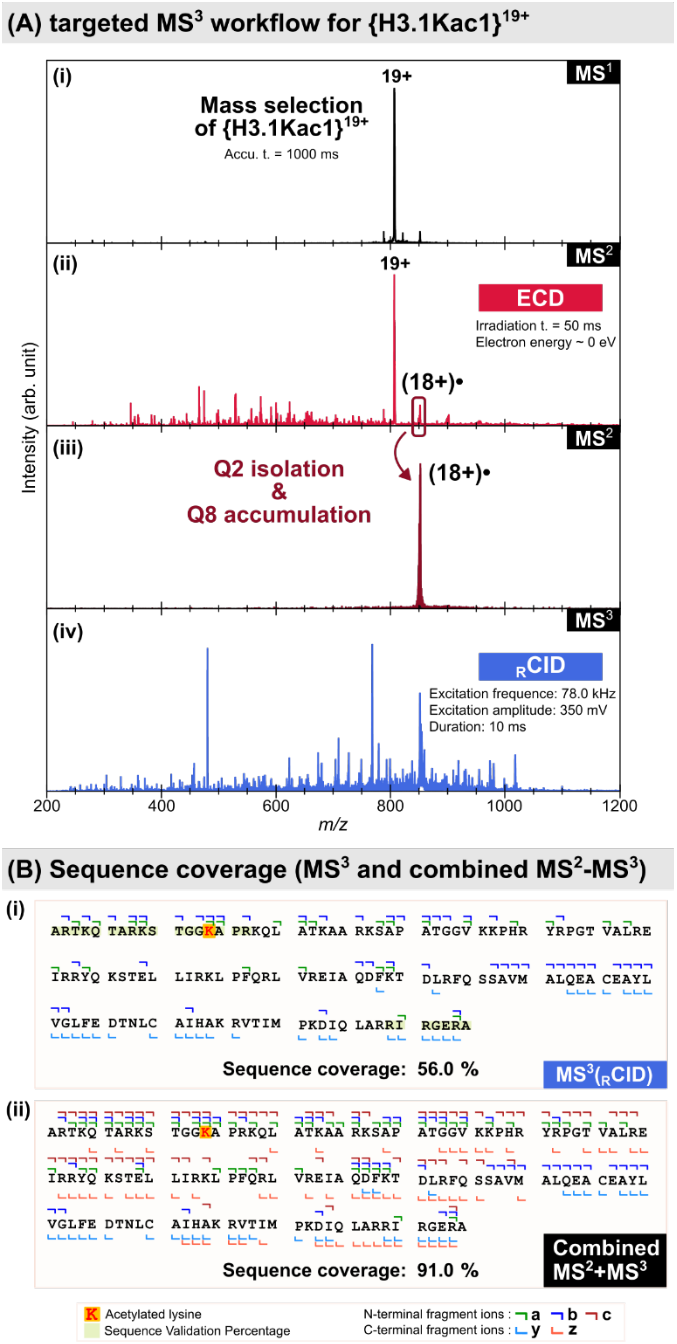

**Figure S1. Multimodal MS$^3$ workflow for {H3.1Kac1} generated by GCN5 (10 min).** (A) Mass spectrum of the mass-selected precursor ion {H3.1Kac}$^{19+}$ (i), subjected to MS$^2$(ECD) to generate charge-reduced species and fragment ions (ii). A charge-reduced species, {H3.1Kac}$^{18+\bullet}$, is isolated and accumulated in Q8 (iii), and subsequently fragmented by MS$^3$($_R$CID) (iv). (B) Fragmentation maps of the identified proteoform [H3.1K14ac] obtained from MS$^3$ dataset (top) and from the combined MS$^2$/MS$^3$ dataset (bottom).

**Table S1. Omniscape ranking of candidate monoacetylated H3.1 proteoforms ({H3Kac1}) across MS² and MS³ experiments following GCN5 treatment, for charge states 19⁺ (MS²: ECD, EID, ECciID, RCID, and MS³: ECD→RCID) and 20⁺ (ECD and RCID).** Proteoforms are ranked according to their MS score. Sequence Validation Percentage (SVP), validated sequence regions, and sequence coverage reported for each assignment. Experimental conditions and number of average spectra (N) are indicated for each dataset.

| MS level | fragmentation | z | Precursor ions | precursor m/z | Proteoform | Rank | MS score | SVP | SVP range | Sequence coverage (%) | Experimental conditions | N |
|---|---|---|---|---|---|---|---|---|---|---|---|---|
| 2 | ECD | 19+ | {H3Kac1} | 806.55 | [H3.1K14ac] | 1 | 36.5 | 81.5 | 1-86, 112-135 | 79.1 | isCID: 40 eV<br>Accumulation time: 1,000 ms<br>ECD irradition time: 50 ms<br>Electron energy ~ 0 eV | 246 |
| | | | | | [H3.1K18ac] | 2 | 34.9 | 27.4 | 1-13, 112-135 | 77.6 | | |
| | | | | | [H3.1K9ac] | 3 | 33.5 | 24.4 | 1-9, 112-135 | 76.1 | | |
| 2 | EID | 19+ | {H3Kac1} | 806.55 | [H3.1K14ac] | 1 | 26.6 | 62.2 | 1-63, 115-135 | 78.4 | Accumulation time: 1,000 ms<br>EID irradiation time: 50 ms<br>Electron energy = 30 eV | 189 |
| | | | | | [H3.1K18ac] | 2 | 25 | 62.2 | 1-63, 115-135 | 76.9 | | |
| | | | | | [H3.1K9ac] | 3 | 24.6 | 22.2 | 1-9, 115-135 | 75.4 | | |
| 2 | ECciD | 19+ | {H3Kac1} | 806.55 | [H3.1K14ac] | 1 | 18.1 | 39.9 | 1-41, 124-135 | 74.6 | isCID: 40 eV<br>Accumulation time: 1,000 ms<br>ECD irradition time: 30 ms<br>CID energy: 31 eV | 242 |
| | | | | | [H3.1K18ac] | 2 | 17.3 | 18.5 | 1-13, 124-135 | 72.4 | | |
| | | | | | [H3.1K9ac] | 3 | 16.6 | 14.8 | 1-8, 124-135 | 70.9 | | |
| 2 | RCID | 19+ | {H3Kac1} | 806.55 | [H3.1K14ac] | 1 | 9.7 | 6.7 | 1-5, 132-135 | 41.8 | Accumulation time: 1,000 ms<br>Excitation frequence: 76.8 kHz<br>Excitation amplitude: 770 mV<br>Duration: 10 ms | 116 |
| | | | | | [H3.1K9ac] | 2 | 9.5 | 6.7 | 1-5, 132-136 | 41 | | |
| | | | | | [H3.1K18ac] | 3 | 9.3 | 6.7 | 1-5, 132-137 | 40.3 | | |
| 2 | ECD | 20+ | {H3Kac1} | 766.27 | [H3.1K14ac] | 1 | 20.8 | 64.4 | 1-64, 113-135 | 68.7 | Accumulation time: 1,000 ms<br>ECD irradition time: 40 ms<br>Collision energy = 55.5 eV | 169 |
| | | | | | [H3.1K18ac] | 2 | 19.3 | 26.7 | 1-13, 113-135 | 67.2 | | |
| | | | | | [H3.1K9ac] | 3 | 19 | 23.7 | 1-9, 113-135 | 65.7 | | |
| 2 | RCID | 20+ | {H3Kac1} | 766.27 | [H3.1K14ac] | 1 | 14.1 | 0 | - | 37.3 | Accumulation time: 1000 ms<br>Excitation frequence: 82.1 kHz<br>Excitation amplitude: 230 mV<br>Duration: 10 ms | 190 |
| | | | | | [H3.1K18ac] | 2 | 13.5 | 0 | - | 35.8 | | |
| | | | | | [H3.1K9ac] | 3 | 13.2 | 0 | - | 35.1 | | |
| 3 | $MS^2$(ECD) | 19+ | {H3Kac1} | 806.55 | | | | | | | Accumulation time: 1,000 ms<br>ECD irradiation time: 20 ms | |
| | $MS^3$(RCID) | 18+• | {H3Kac1} | 851.37 | [H3.1K14ac] | 1 | 16 | 17.8 | 1-17, 129-135 | 56 | Excitation frequence: 78.0 kHz<br>Excitation amplitude: 350 mV<br>Duration: 10 ms | 366 |
| | | | | | [H3.1K9ac] | 2 | 15.3 | 11.1 | 1-8, 129-135 | 53.7 | | |
| | | | | | [H3.1K18ac] | 3 | 15.2 | 14.8 | 1-13, 129-135 | 53.7 | | |

**Table S2. Omniscape ranking of the top three candidate monoacetylated H3.1 proteoforms assigned to the precursor ion {H3Kac1}$^{19+}$ generated after 30 min PCAF treatment.** MS$^2$ datasets acquired by ECD, EID, and $_{\mathrm{R}}$CID are shown. Proteoforms are ranked by MS score, and the corresponding Sequence Validation Percentage (SVP), validated sequence region, and sequence coverage are reported for each assignment. Experimental conditions and the number of average spectra (N) are indicated for each dataset.

| MS level | fragmentation | z | Precursor ions | precursor m/z | Proteoform | Rank | MS score | SVP | SVP range | Sequence coverage (%) | Experimental conditions | N |
|---|---|---|---|---|---|---|---|---|---|---|---|---|
| 2 | ECD | 19+ | {H3Kac1} | 806.55 | [H3.1K14ac] | 1 | 28.7 | 67.4 | 1-64, 109-135 | 81.3 | isCID: 30 eV<br>Accumulation time:<br>400 ms<br>ECD irradition time:<br>60 ms<br>Electron energy ~ 0 eV | 432 |
| | | | | | [H3.1K18ac] | 2 | 27.1 | 29.6 | 1-13. 109-135 | 78.4 | | |
| | | | | | [H3.1K9ac] | 3 | 26.7 | 27.4 | 1-10, 109-135 | 79.9 | | |
| 2 | EID | 19+ | {H3Kac1} | 806.55 | [H3.1K14ac] | 1 | 13.3 | 45.2 | 1-41, 116-135 | 61.2 | isCID: 30 eV<br>Accumulation time:<br>400 ms<br>EID irradition time: 40 ms<br>Electron energy ~ 35 eV | 921 |
| | | | | | [H3.1K9ac] | 2 | 12.5 | 45.9 | 1-41, 115-135 | 61.2 | | |
| | | | | | [H3.1K18ac] | 3 | 12.0 | 24.4 | 1-13, 116-135 | 59 | | |
| 2 | $_{\mathrm{R}}$CID | 19+ | {H3Kac1} | 806.55 | [H3.1K14ac] | 1 | 23.5 | 8.9 | 1-5, 129-135 | 48.5 | isCID : 30 eV<br>Accumulation time:<br>150 ms<br>Excitation frequence:<br>69.7 kHz<br>Excitation amplitude:<br>335 mV<br>Duration: 10 ms | 790 |
| | | | | | [H3.1K9ac] | 2 | 23.2 | 8.9 | 1-5, 129-135 | 47.8 | | |
| | | | | | [H3.1K18ac] | 3 | 22.4 | 8.9 | 1-5, 129-135 | 46.3 | | |

**Table S3. Omniscape ranking of the top four candidate monoacetylated H4 proteoforms assigned to {H4Kac1}$^{13+}$ after 30 min PCAF treatment.** MS$^2$ datasets acquired by ECD, EID, ECciD, and $_{R}$CID. MS$^3$ data derived from $_{R}$CID of {H4Kac1}$^{13+}$ and subsequent ECD of the isolated monoacetylated $b_{52}^{7+}$ fragment ion are shown. Proteoforms are ranked by MS score, with the corresponding Sequence Validation Percentage (SVP), validated sequence region, and sequence coverage reported for each assignment. Experimental conditions and the number of average spectra (N) are indicated for each dataset.

| MS level | fragmentation | z | Precursor ions | precursor m/z | Proteoform | Rank | MS score | SVP | SVP range | Sequence coverage (%) | Experimental conditions | N |
|---|---|---|---|---|---|---|---|---|---|---|---|---|
| 2 | ECD | 13+ | {H4Kac1} | 868.03 | [H4K8ac] | 1 | 63 | 97.1 | 1-64, 68-102 | 91.1 | isCID: 40 eV<br>Accumulation time: 100 ms<br>ECD irradition time: 80 ms<br>Electron energy ~ 0 eV | 572 |
| | | | | | [H4K12ac] | 2 | 62 | 97.1 | 1-64, 68-102 | 91.1 | | |
| | | | | | [H4K16ac] | 3 | 59.9 | 97.1 | 1-64, 68-102 | 91.1 | | |
| | | | | | [H4K5ac] | 4 | 59.7 | 38.2 | 1-4, 68-102 | 88.1 | | |
| 2 | EID | 13+ | { H4Kac1} | 868.03 | [H4K8ac] | 1 | 42.3 | 66.7 | 1-63, 98-102 | 89.1 | isCID: 40 eV<br>Accumulation time: 80 ms<br>EID irradition time: 80 ms<br>Electron energy ~ 35 eV | 692 |
| | | | | | [H4K12ac] | 2 | 41.1 | 66.7 | 1-63, 98-102 | 89.1 | | |
| | | | | | [H4K5ac] | 3 | 40.4 | 8.8 | 1-4, 98-102 | 86.1 | | |
| | | | | | [H4K16ac] | 4 | 39.7 | 66 | 1-63, 98-102 | 88.1 | | |
| 2 | ECciD | 13+ | { H4Kac1} | 868.03 | [H4K8ac] | 1 | 20.6 | 35.73 | 1-31, 98-102 | 77.2 | isCID : 40 eV<br>Accumulation time: 100 ms<br>ECD irradition time: 80 ms<br>CID energy: 30 eV | 686 |
| | | | | | [H4K12ac] | 2 | 19.8 | 35.3 | 1-31, 98-102 | 77.2 | | |
| | | | | | [H4K16ac] | 3 | 19.2 | 16.7 | 1-12, 98-102 | 76.2 | | |
| | | | | | [H4K5ac] | 4 | 18.6 | 8.8 | 1-4, 98-102 | 74.3 | | |
| 2 | $_{R}$CID | 13+ | {H4Kac1} | 868.03 | [H4K8ac] | 1 | 19.9 | 23.5 | 1-19, 98-102 | 58.4 | isCID : 40 eV<br>Accumulation time: 30 ms<br>Excitation frequence: 67.8 kHz<br>Excitation amplitude: 300 mV<br>Duration: 10 ms | 1536 |
| | | | | | [H4K12ac] | 2 | 19.4 | 23.5 | 1-19, 98-102 | 58.4 | | |
| | | | | | [H4K5ac] | 3 | 18.5 | 8.8 | 1-4, 98-102 | 55.4 | | |
| | | | | | [H4K16ac] | 4 | 18.4 | 16.7 | 1-12, 98-102 | 56.4 | | |
| 3 | MS$^2$($_{R}$CID) | 13+ | {H4Kac1} | 868.03 | | | | | | | isCID : 40 eV<br>Accumulation time: 100 ms<br>Excitation frequence: 67.8 kHz<br>Excitation amplitude: 300 mV<br>Duration: 10 ms | |
| | MS$^3$(ECD) | $b_{52}^{7+}$ | {H4$_{[1-52]}$Kac1} | 803.19 | [H4$_{[1-52]}$K8ac] | 1 | 28.1 | 80.8 | 1-40, 51-52 | 82.4 | ECD irradition time: 80 ms<br>Electron energy ~ 0 eV | 360 |
| | | | | | [H4$_{[1-52]}$K12ac] | 2 | 26.1 | 21.2 | 1-9, 51-52 | 78.4 | | |
| | | | | | [H4$_{[1-52]}$K16ac] | 3 | 25.0 | 21.2 | 1-9, 51-52 | 82.4 | | |
| | | | | | [H4$_{[1-52]}$K5ac] | 4 | 24.9 | 11.5 | 1-4, 51-52 | 78.4 | | |

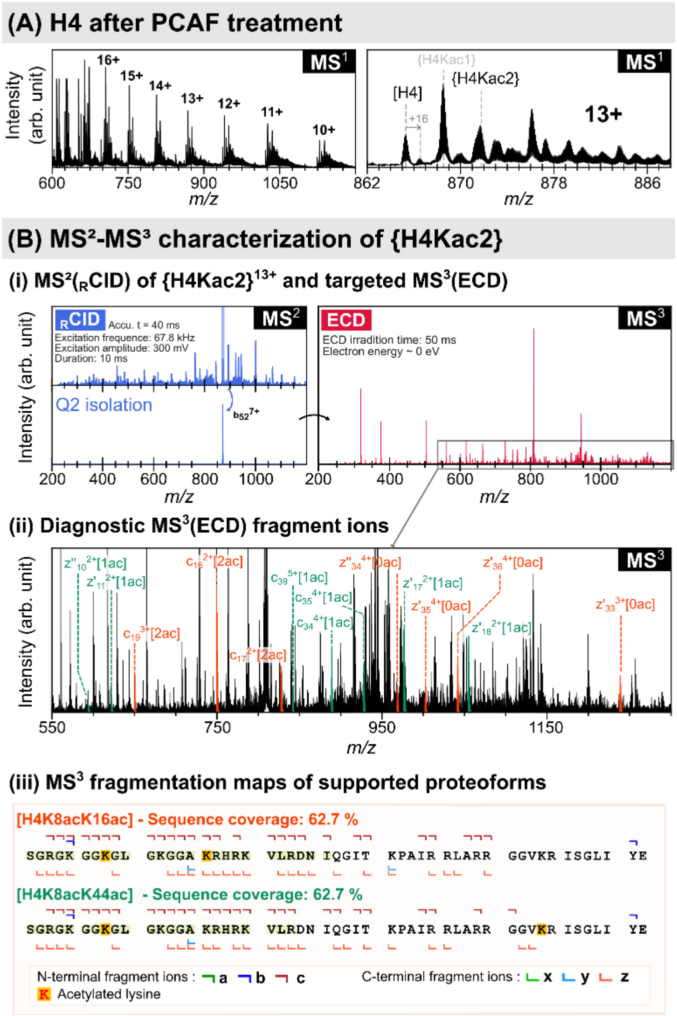

**Figure S2. MS$^2$/ targeted MS$^3$ analysis of the doubly acetylated histone H4 species {H4Kac2}$^{13+}$ generated by PCAF treatment.** (A) MS$^1$ spectrum of H4 after 30 min PCAF treatment, with a zoomed view of charge state 13+ region. (B) MS$^2$ and targeted MS$^3$ workflow applied to {H4Kac2}$^{13+}$. (i) MS$^2$(RCID) spectrum (top left). Isolation of $b_{52}^{7+}$ fragment, carrying two acetylations, generated by RCID was isolated in Q2 and accumulated in Q8 (bottom left), and subsequent MS$^3$(ECD) spectrum (right). (ii) Expanded view of selected MS$^3$ fragment ions supporting the assignments [H4K8acK16ac] and [H4K8acK44ac]. (iii) MS$^3$ fragmentation maps for [H4K8acK16ac] and [H4K8acK44ac].

The {H4Kac2}$^{13+}$ species generated by PCAF incubation presented a more complex analytical case than monoacetylated species because it was detected at lower intensity and partially overlapped with background isobaric/isomeric contaminant (Figure S2A, right panel). Consequently, MS score were lower than those obtained for {H4Kac1} (a decrease of ~30 % for ECD). Both MS$^2$(ECD) and MS$^2$(RCID) consistently rank [H4K8acK44ac] as the top assignment, with closely related candidates ([H4K8acK31ac], [H4K8acK16ac], [H4K8K20ac], [H4K16K44ac]) within a narrow MS score range. However, no clear evidence supports acetylation beyond K44. To refine the localization of acetyl marks, MS$^3$(ECD) was performed on diacetylated $b_{52}^{7+}$ fragment generated by RCID. In this spectrum, [H4K8acK31] and [H4K8acK44] were the two highest scoring proteoforms (MS score 10.5 and 9.9, respectively). However, they did not yield the highest SVP (44.2 %), with coverage restricted to fragments spanning residues up to 23. In contrast, [H4K8acK20ac] and [H4K8acK16ac] displayed slightly lower MS scores (9.5 and 8.7, respectively) but achieved higher SVP values (50%), with fragment coverage extending to residue 26. These were the only proteoform for which both acetylation sites felt within the SVP-supported region.

Inspection of the MS$^3$ fragment ions provided site constraints. The $c_5$-$c_7$ fragments were detected only unmodified, whereas $c_8$-$c_{11}$ fragments were detected only with one acetylation, establishing K8 as a conserved acetylation site within the {H4Kac2} population. The absence of doubly acetylated fragments before to residue

15 excluded [H4K8acK12ac]. Consequently, the additional acetylation was localized within the distal N-terminal region (K16-K44). Direct support of [H4K8acK16ac] was provided by doubly acetylated fragments $c_{16\text{-}17}$ and $c_{19}$ with unmodified C-terminal fragment ions $z_{33}$-$z_{36}$. From the C-terminal direction, sparse low-intensity z-type ($z_{10\text{-}11}$, $z_{17\text{-}18}$) bearing a single acetylation, alongside monoacetylated N-terminal fragments unmodified $c_{34\text{-}35,\ 39}$ were compatible with [H4K8acK44ac]. In the absence of site-specific diagnostic fragments, intermediate lysines cannot be uniquely localized. Thus, K8ac was firmly established within the {H4Kac2} species, while [H4K8acK16ac] and [H4K8acK44ac] were supported by site-specific ions, intermediate residues such as K20 and K31 remained plausible but could not be uniquely assigned under these fragmentation conditions.

**Table S4. Omniscape ranking of the top four candidate diacetylated H4 proteoforms assigned to $\{H4Kac2\}^{13+}$ precursor generated after 30 min PCAF treatment.** $MS^2$ datasets acquired by ECD, EID, and $_R$CID are shown together with targeted $MS^3$(ECD) analysis of the isolated diacetylated $b_{52}^{7+}$ fragment ($\{H4_{[1-52]}Kac2\}^{7+}$) generated by $_R$CID. Proteoforms are ranked by MS score, with the corresponding Sequence Validation Percentage (SVP), validated sequence region, and sequence coverage reported for each assignment. Experimental conditions and the number of average spectra (N) are indicated for each dataset.

| MS level | fragmentation | z | Precursor ions | precursor m/z | Proteoform | Rank | MS score | SVP | SVP range | Sequence coverage (%) | Experimental conditions | N |
|---|---|---|---|---|---|---|---|---|---|---|---|---|
| 2 | ECD | 13+ | {H4Kac2} | 871.26 | [H4K8acK44ac] | 1 | 42.8 | 73.5 | 1-60, 88-102 | 85.1 | isCID: 40 eV<br>Accumulation time: 100 ms<br>ECD irradition time: 100 ms<br>Electron energy ~ 0 eV | 908 |
| | | | | | [H4K12acK44ac] | 2 | 42.3 | 73.5 | 1-64, 88-102 | 85.1 | | |
| | | | | | [H4K16acK44ac] | 3 | 41.0 | 73.5 | 1-64, 88-102 | 85.1 | | |
| | | | | | [H4K8acK31ac] | 4 | 40.4 | 73.5 | 1-64, 88-102 | 84.2 | | |
| 2 | $_R$CID | 13+ | {H4Kac2} | 871.26 | [H4K8acK20ac] | 1 | 13.4 | 25.5 | 1-19, 95-102 | 56.4 | isCID : 40 eV<br>Accumulation time: 30 ms<br>Excitation frequence: 67.7 kHz<br>Excitation amplitude: 300 mV<br>Duration: 10 ms | 945 |
| | | | | | [H4K12acK20ac] | 2 | 13.0 | 25.5 | 1-19, 95-102 | 56.4 | | |
| | | | | | [H4K5acK20ac] | 3 | 12.8 | 10.8 | 1-4, 95-102 | 54.5 | | |
| | | | | | [H4K8acK16ac] | 4 | 12.8 | 21.6 | 1-15, 95-102 | 54.5 | | |
| 3 | $MS^2$($_R$CID) | 13+ | {H4Kac2} | 871.26 | | | | | | | isCID : 40 eV<br>Accumulation time: 40 ms<br>Excitation frequence: 67.7 kHz<br>Excitation amplitude: 300 mV<br>Duration: 10 ms | |
| | $MS^3$(ECD) | $b_{52}^{7+}$ | $\{H4_{[1-52]}Kac2\}$ | 809.19 | [H4K8acK31ac] | 1 | 10.5 | 44.2 | 1-23 | 64.7 | ECD irradition time: 50 ms<br>Electron energy ~ 0 eV | 225 |
| | | | | | [H4K8acK44ac] | 2 | 9.9 | 44.2 | 1-23 | 62.7 | | |
| | | | | | [H4K12acK31ac] | 3 | 9.5 | 13.5 | 1-7 | 60.8 | | |
| | | | | | [H4K8acK20ac] | 4 | 9.5 | 50 | 1-26 | 62.7 | | |

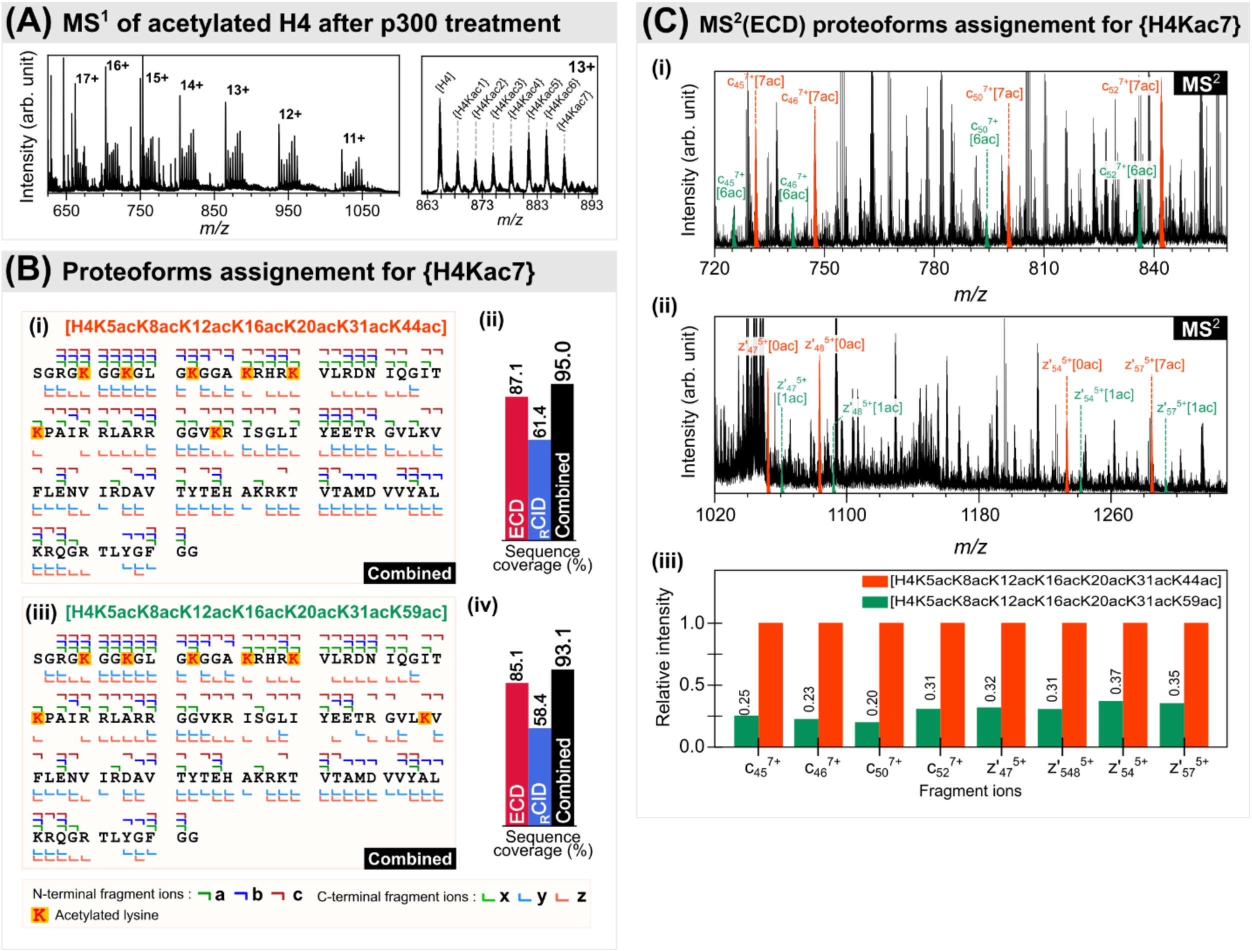

**Figure S3. Assignment of {H4Kac7} proteoforms generated by p300 after 60 minutes of reaction.** (A) $MS^1$ spectrum and zoomed view of the charge-state $13^+$ region containing {H4Kac7}. (B) Combined fragmentation maps and sequence coverage histograms for the two candidate proteoforms identified from ECD and RCID: (i, ii) [H4K5acK8acK12acK16acK20acK31acK44ac] and (iii, iv) [H4K5acK8acK12acK16acK20acK31acK59ac]. (C) ECD spectral regions showing common diagnostic c-type ions (i) and complementary z-type ions (ii) supporting both positional isomers, shown in orange for the K44ac proteoform and green for the K59ac proteoform; (iii) relative intensities of the seletecd diagnostic fragment ions.

**H4 species carrying seven acetylations generated by p300 after 1 hour.** $MS^2$(ECD) and $MS^2$(RCID) were performed on $\{H4Kac7\}^{13+}$ precursor. Both activation methods ranked [H4K5acK8acK12acK16acK20acK31acK44ac] as the top-scoring proteoform. ECD provided 87.1% sequence coverage and continuous SVP up to residue 64, with an MS score approximately 10% higher than the alternative assignment [H4K5acK8acK12acK16acK20acK31acK59ac], for which SVP terminated at residue 56. The ECD data contained c-type fragments spanning residues 45-58 with either six or seven acetylations together with complementary z-type fragments carrying one or zero acetylations, supporting the presence of an acetylation downstream of K31. Alternative configurations lacking K31ac were disfavored because SVP terminated near residue 30. Notably, no C-terminal fragments validating acetylation beyond residue ~59 were consistently observed, resulting in a gap in sequence coverage across residues 57–91 and effectively excluding more distal lysine sites from consideration for the seventh acetylation. RCID provided 61.4 % sequence coverage for [H4K5acK8acK12acK16acK20acK31acK44ac] proteoform but generated limited fragment evidence across residues 30-50, restricting its ability to localize the terminal acetylations. Thus, under these conditions, RCID confirmed the acetylations at K5-K20 but did not enable unambiguous localization of the sixth and seventh acetylation beyond K20. Taken together, these constraints restricted the hepta-acetylated population to two positional isomers, differing by acetylation at either K44 or K59. Combined ECD and RCID sequence coverage led to 95.0% and 93.1% sequence coverage of for

[H4K5acK8acK12acK16acK20acK31acK44ac] and [H4K5acK8acK12acK16acK20acK31acK59ac], respectively.

The relative abundance of these two isomers differing at K44ac/K59ac was estimated from the most abundant diagnostic c-ions ($c_{45}^{7+}$, $c_{46}^{7+}$, $c_{50}^{7+}$, $c_{52}^{7+}$), and complementary z ions ($z'_{47}{}^{5+}$, $z'_{48}{}^{5+}$, $z'_{54}{}^{5+}$, $z'_{57}{}^{5+}$) extracted from the same ECD experiments and compared at identical charge states using their maximum ion intensities. This analysis estimated approximately 75-80% [H4K5acK8acK12acK16acK20acK31acK44ac] and 20-25% [H4K5acK8acK12acK16acK20acK31acK59ac] within the {H4Kac7} population.

**Table S5. Omniscape ranking of the top four candidate hexa- and hepta-acetylated H4 proteoforms assigned to {H4Kac6}$^{13+}$ and {H4Kac7}$^{13+}$ generated after p300 treatment.** MS$^2$ datasets acquired by ECD, EID, ECciD, and $_{R}$CID are shown for {H4Kac6}$^{13+}$, and ECD and $_{R}$CID are also shown for {H4Kac7}$^{13+}$. Targeted MS$^3$(ECD) data obtained from $_{R}$CID generation and isolation of the $c_{22}^{3+}$ fragment ion {H4$_{[1\text{-}22]}$Kac5}$^{3+}$ are also reported. Proteoforms are ranked by MS score, with the corresponding Sequence Validation Percentage (SVP), validated sequence region, and sequence coverage reported for each assignment. Experimental conditions and the number of average spectra (N) are indicated for each dataset.

| MS level | fragmentation | z | Precursor ions | precursor m/z | Proteoform | Rank | MS score | SVP | SVP range | Sequence coverage (%) | Experimental conditions | N |
|---|---|---|---|---|---|---|---|---|---|---|---|---|
| 2 | ECD | 13+ | {H4Kac6} | 884.18 | [H4K5acK8acK12acK16acK20acK44ac] | 1 | 52.5 | 54.9 | 1-41, 88-102 | 89.1 | isCID: 40 eV<br>Accumulation time: 250 ms<br>ECD irradition time: 100 ms<br>Electron energy ~ 0 eV | 676 |
| | | | | | [H4K5acK8acK12acK16acK20acK31ac] | 2 | 49.2 | 43.1 | 1-29, 88-102 | 87.1 | | |
| | | | | | [H4K5acK8acK12acK16acK31acK44ac] | 3 | 46.9 | 41.2 | 1-27, 88-102 | 87.1 | | |
| | | | | | [H4K5acK8acK12acK20acK31acK44ac] | 4 | 45.6 | 41.2 | 1-27, 88-102 | 87.1 | | |
| 2 | EID | 13+ | {H4Kac6} | 884.18 | [H4K5acK8acK12acK16acK20acK44ac] | 1 | 35.3 | 50.0 | 1-46, 98-102 | 90.1 | isCID: 40 eV<br>Accumulation time: 300 ms<br>EID irradition time: 100 ms<br>Electron energy ~ 35 eV | 598 |
| | | | | | [H4K5acK8acK12acK16acK20acK31ac] | 2 | 33.5 | 35.3 | 1-31, 98-102 | 89.1 | | |
| | | | | | [H4K5acK8acK12acK16acK20acK59ac] | 3 | 31.3 | 47.1 | 1-43, 98-102 | 88.1 | | |
| | | | | | [H4K5acK8acK12acK16acK31acK44ac] | 4 | 30.3 | 30.4 | 1-26, 98-102 | 87.1 | | |
| 2 | ECciD | 13+ | {H4Kac6} | 884.18 | [H4K5acK8acK12acK16acK20acK44ac] | 1 | 40.8 | 34.3 | 1-30, 98-102 | 87.1 | isCID : 40 eV<br>Accumulation time: 400 ms<br>ECD irradition time:80 ms<br>CID energy: 30 eV | 770 |
| | | | | | [H4K5acK8acK12acK16acK20acK31ac] | 2 | 37.8 | 34.3 | 1-30, 98-102 | 86.1 | | |
| | | | | | [H4K5acK8acK12acK16acK31acK44ac] | 3 | 34.4 | 23.5 | 1-19, 98-102 | 82.2 | | |
| | | | | | [H4K5acK8acK12acK20acK31acK44ac] | 4 | 33.2 | 23.5 | 1-19, 98-102 | 81.2 | | |
| 2 | $_{R}$CID | 13+ | {H4Kac6} | 884.18 | [H4K5acK8acK12acK16acK20acK44ac] | 1 | 20.6 | 16.7 | 1-11, 97-102 | 52.5 | isCID : 40 eV<br>Accumulation time: 200 ms<br>Excitation frequence: 67.2 kHz<br>Excitation amplitude: 350 mV<br>Duration: 10 ms | 670 |
| | | | | | [H4K5acK8acK12acK16acK31acK44ac] | 2 | 20.2 | 16.7 | 1-11, 97-102 | 51.5 | | |
| | | | | | [H4K5acK8acK12acK16acK20acK31ac] | 3 | 19.7 | 16.7 | 1-11, 97-102 | 50.5 | | |
| | | | | | [H4K5acK8acK12acK20acK31acK44ac] | 4 | 19.7 | 16.7 | 1-11, 97-102 | 50.5 | | |
| 2 | ECD | 13+ | {H4Kac7} | 887.48 | [H4K5acK8acK12acK16acK20acK31acK44ac] | 1 | 50.6 | 62.7 | 1-64 | 87.1 | isCID: 40 eV<br>Accumulation time: 500 ms<br>ECD irradition time: 80 ms<br>Electron energy ~ 0 eV | 486 |
| | | | | | [H4K5acK8acK12acK16acK20acK31acK59ac] | 2 | 46.0 | 54.9 | 1-56 | 85.1 | | |
| | | | | | [H4K5acK8acK12acK16acK20acK44acK59ac] | 3 | 40.1 | 29.4 | 1-30 | 82.2 | | |
| | | | | | [H4K5acK8acK12acK16acK20acK31acK77ac] | 4 | 38.8 | 54.9 | 1-56 | 80.2 | | |
| | $_{R}$CID | 13+ | {H4Kac7} | 887.42 | [H4K5acK8acK12acK16acK20acK31acK44ac] | 1 | 26.2 | 23.5 | 1-19, 98-102 | 61.4 | isCID : 40 eV<br>Accumulation time: 100 ms<br>Excitation frequence: 67.1 kHz<br>Excitation amplitude: 350 mV<br>Duration: 10 ms | 732 |
| | | | | | H4K5acK8acK12acK16acK20acK44acK59ac] | 2 | 23.4 | 20.6 | 1-16, 98-102 | 59.4 | | |
| | | | | | H4K5acK8acK12acK16acK20acK31acK59ac] | 3 | 23.3 | 20.6 | 1-16, 98-102 | 58.4 | | |
| | | | | | H4K5acK8acK12acK16acK31acK44acK59ac] | 4 | 21.5 | 20.6 | 1-16, 98-102 | 55.4 | | |
| 3 | MS$^2$($_{R}$CID) | 13+ | {H4Kac6} | 887.42 | | | | | | | isCID : 40 eV<br>Accumulation time: 100 ms<br>Excitation frequence: 67.1 kHz<br>Excitation amplitude: 350 mV<br>Duration: 10 ms | |
| | MS$^3$(ECD) | $C_{22}^{3+}$ | {H4$_{[1\text{-}22]}$Kac5} | 805.13 | [H4K5acK8acK12acK16acK20ac] | 1 | 17.0 | 100 | 1-22 | 95.2 | ECD irradition time: 100 ms<br>Electron energy ~ 0 eV | 183 |

### Comparison of mono- and bi-exponential models for sequence coverage accumulation of [H3.1K14ac] generated by PCAF treatment of H3.1 (30 min)

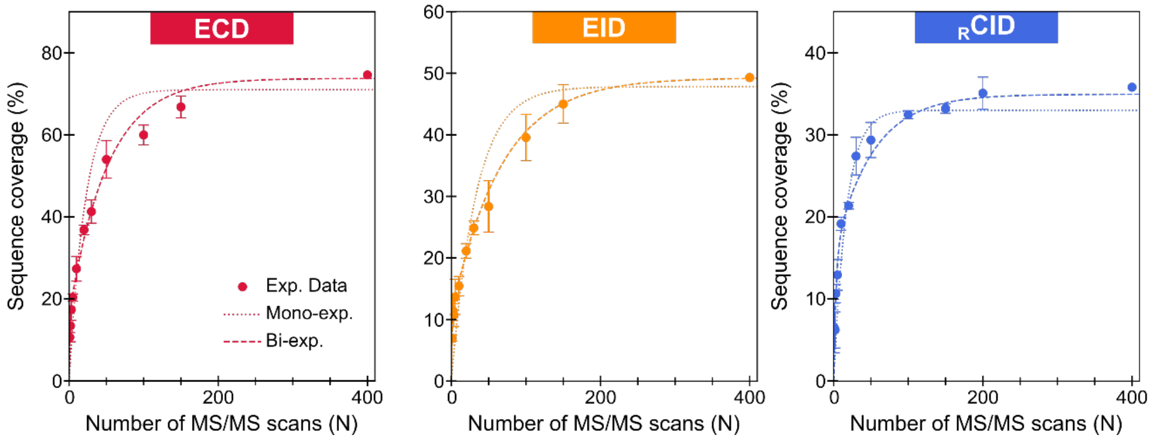

**Figure S4. Mono- and bi-exponential fitting of sequence coverage accumulation curves for [H3.1K14ac] generated after 30 min PCAF treatment of H3.1.** Experimental sequence coverage values obtained by ECD (red), EID (orange), and RCID (blue) are plotted as a function of the number of accumulated $MS^2$ scans. Adjusted $R^2$ values for each mono-exponential and bi-exponential are: ECD, 0.91569 and 0.99954; EID, 0.8991 and 0.9998; RCID, 0.9311 and 0.9994, respectively.

#### Adjusted coefficient of determination

The adjusted coefficient of determination $R^2$ is defined as:

$$R^2 = 1 - \frac{\frac{1}{n-p-1}\sum_{i=1}^{n}\left(SC_i - \widehat{SC}_i\right)^2}{\frac{1}{n-1}\sum_{i=1}^{n}\left(SC_i - \overline{SC}\right)^2}$$

Where $C_i$ are the experimental sequence coverage values, $\widehat{SC}_i$ are the model-predicted values, $\overline{SC}$ is the mean experimental sequence coverage, n is the number of data points, p is the number of fitted parameters (p=2 for the mono-exponential model and p=4 for the bi-exponential model). The adjusted $R^2$ allows for direct comparison between models of different complexity by taking into account the number of parameters involved in each fit.

#### Mono-exponential saturation model

$$C(N) = C_\infty(1 - e^{-kN})$$

Where $C(N)$ is the sequence coverage obtained from N MS/MS spectra, $C_\infty$ is the asymptotic coverage plateau, $k$ describes the rate of accumulation of new backbone cleavages.

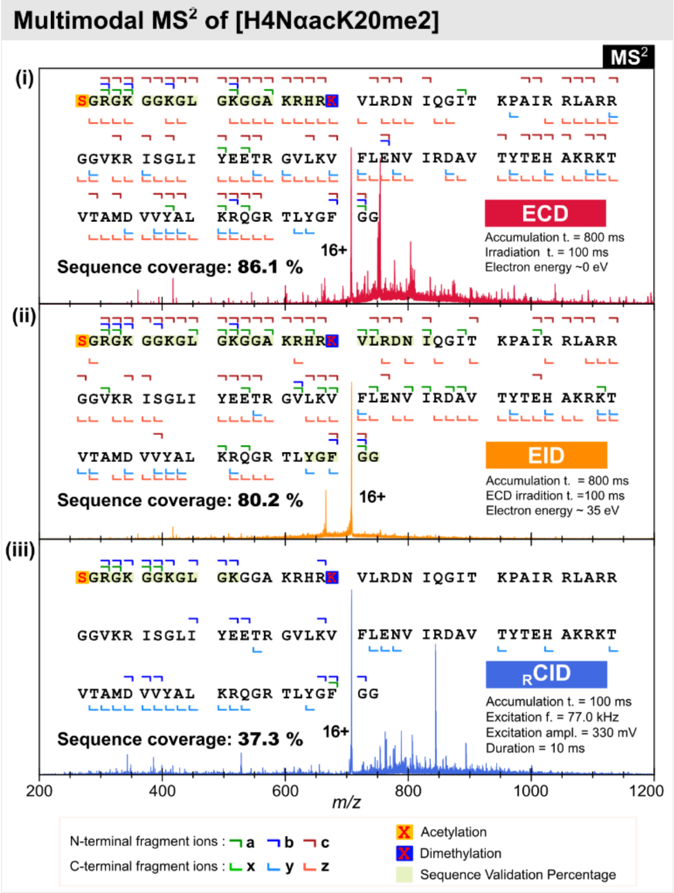

**Figure S5. Multimodal MS² fragmentation analysis of the endogenous histone H4 proteoform [H4NαacK20me2].** Representative ECD (i), EID (ii) and $_{R}$CID (iii) spectra of the $16^+$ precursor at *m/z* 707 are shown with the corresponding fragment assignments along the H4 sequence.

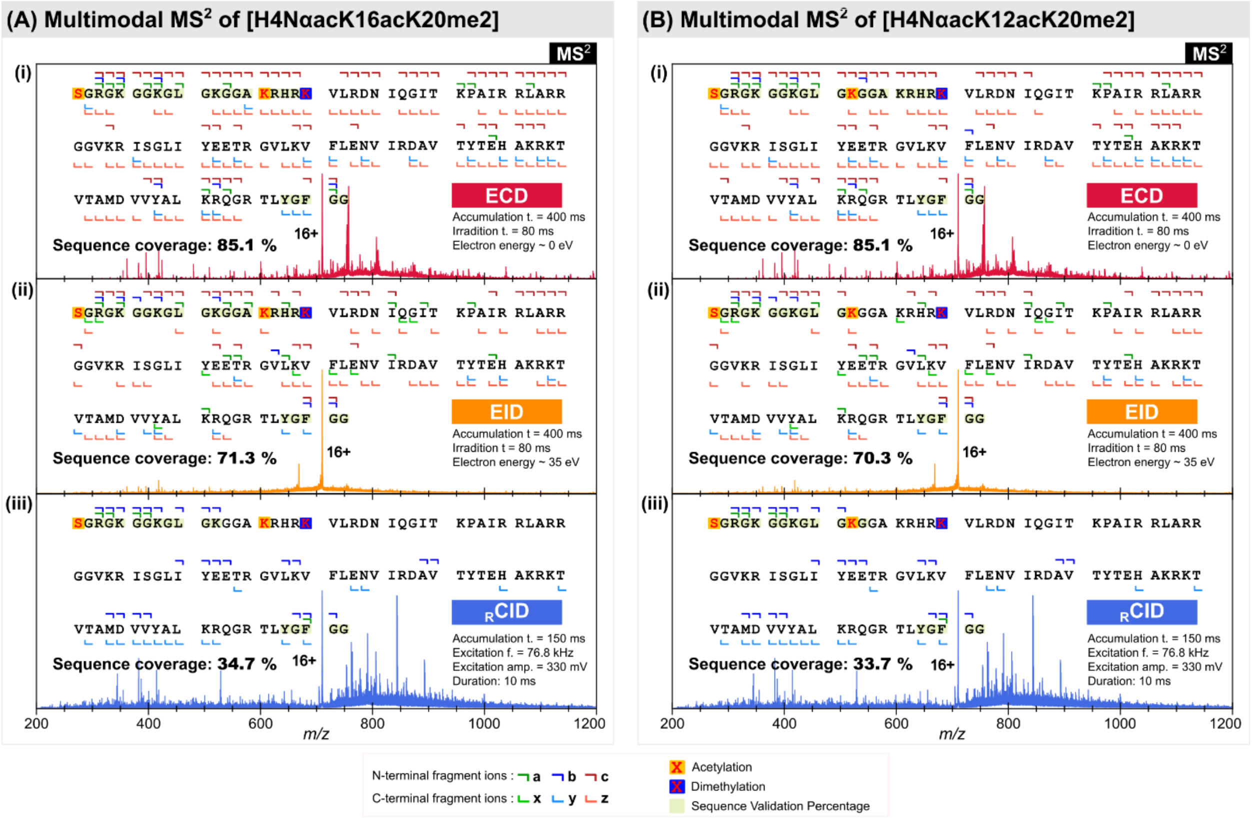

**Figure S6. Multimodal MS² fragmentation analysis of endogenous histone H4 proteoforms at precursor *m/z* 710 (16+).** Representative ECD (i), EID (ii), and $_{R}$CID (iii) spectra are shown for the candidate proteoforms [H4NαacK16acK20me2] (A) and [H4NαacK12acK20me2] (B), together with the corresponding fragment assignments along the H4 sequence.

**(i) Endogenous H4 ECD dataset, precursor *m/z* 707**

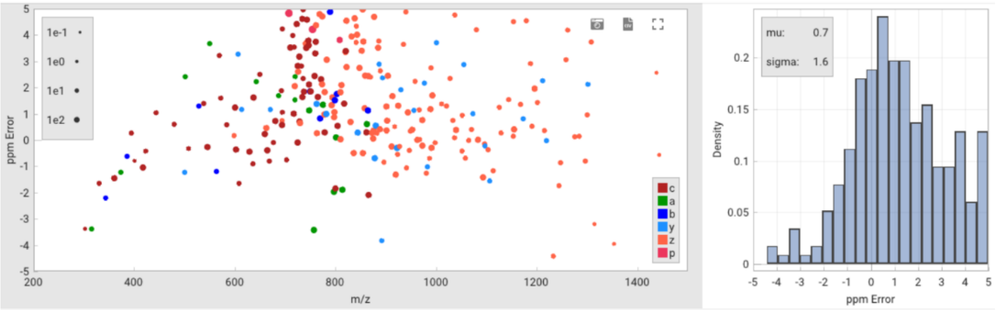

**(ii) Endogenous H4 ECD dataset, precursor *m/z* 710**

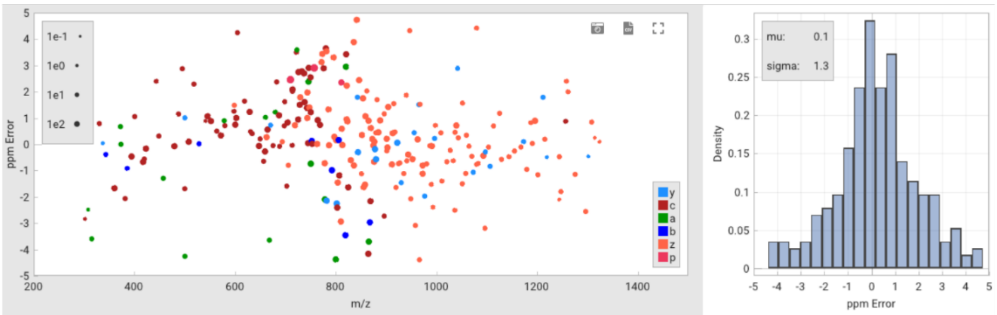

**Figure S7. Fragment mass error plots for the recalibrated ECD datasets of endogenous histone H4.** Mass errors are shown for fragments ions assigned to the precursor ions at *m/z* 707 (i) and 710 (ii). For each dataset, ppm errors are plotted as a function of fragment m/z (left) and as error histograms (right).

**Table S6. Omniscape ranking of the top three candidate endogenous histone H4 proteoforms corresponding to the precursor ions {H4ac1Me2}$^{16+}$ (*m/z* 707) and {H4Ac2Me2}$^{16+}$ (*m/z* 710) detected in liver cell extracts.** $MS^2$ datasets acquired by ECD, EID and RCID are shown. Proteoforms are ranked by MS score, the corresponding Sequence Validation Percentage (SVP), validated sequence region, and sequence coverage are reported for each assignment. Experimental conditions and the number of average spectra (N) are indicated for each dataset.

| MS level | fragmentation | z | Precursor ions | precursor m/z | Proteoform | Rank | MS score | SVP | SVP range | Sequence coverage (%) | Experimental conditions | N |
|---|---|---|---|---|---|---|---|---|---|---|---|---|
| 2 | ECD | 16 | {H4Ac1Me2} | 707.21 | [H4NαacK20me2] | 1 | 39.1 | 18.6 | 1-19 | 86.1 | isCID: 30 eV<br>Accumulation time: 800 ms<br>ECD irradition time: 100 ms<br>Electron energy ~ 0 eV | 311 |
| | | | | | [H4K5acK20me2] | 2 | 38.0 | 0.0 | - | 85.1 | | |
| | | | | | [H4K16acK20me2] | 3 | 36.7 | 14.7 | 1-15 | 82.2 | | |
| 2 | EID | 16 | {H4Ac1Me2} | 707.21 | [H4NαacK20me2] | 1 | 28.3 | 30.4 | 1-26, 98-102 | 80.2 | isCID: 30 eV<br>Accumulation time: 800 ms<br>EID irradition time: 100 ms<br>Electron energy ~ 35 eV | 313 |
| | | | | | [H4K5acK20me2] | 2 | 28.3 | 30.4 | 1-26, 98-102 | 79.2 | | |
| | | | | | [H4K8acK20me2] | 3 | 27.9 | 7.8 | 1-3, 98-102 | 78.2 | | |
| 2 | RCID | 16 | {H4Ac1Me2} | 707.21 | [H4NαacK20me2] | 1 | 9.6 | 11.8 | 1-12 | 35.6 | isCID : 40 eV<br>Accumulation time: 100 ms<br>Excitation frequence: 77.0 kHz<br>Excitation amplitude: 330 mV<br>Duration: 10 ms | 1096 |
| | | | | | [H4NαacK16me2] | 2 | 9.3 | 11.8 | 1-12 | 34.7 | | |
| | | | | | [H4NαacK16me1K20me1] | 3 | 9.3 | 11.8 | 1-12 | 34.7 | | |
| 2 | ECD | 16 | {H4Ac2Me2} | 709.84 | [H4NαacK16acK20me2] | 1 | 35.4 | 23.5 | 1-19, 98-102 | 85.1 | isCID: 30 eV<br>Accumulation time: 400 ms<br>ECD irradition time: 80 ms<br>Electron energy ~ 0 eV | 478 |
| | | | | | [H4K5acK16acK20me2] | 2 | 35.2 | 4.9 | 98-102 | 84.2 | | |
| | | | | | [H4K5acK12acK20me2] | 3 | 34.1 | 23.5 | 1-19, 98-102 | 85.1 | | |
| 2 | EID | 16 | {H4Ac2Me2} | 709.84 | [H4K5acK16acK20me2] | 1 | 16.6 | 4.9 | 98-102 | 71.3 | isCID: 30 eV<br>Accumulation time: 400 ms<br>EID irradition time: 80 ms<br>Electron energy ~ 35 eV | 356 |
| | | | | | [H4NαacK16acK20me2] | 2 | 16.5 | 23.5 | 1-19, 98-102 | 73.3 | | |
| | | | | | [H4K8acK16acK20me2] | 3 | 16.1 | 4.9 | 98-102 | 69.3 | | |
| 2 | RCID | 16 | {H4Ac2Me2} | 709.84 | [H4NαacK16me2K20ac] | 1 | 5.6 | 16.7 | 1-12, 98-102 | 34.7 | isCID : 30 eV<br>Accumulation time: 150 ms<br>Excitation frequence: 76.8 kHz<br>Excitation amplitude: 330 mV<br>Duration: 10 ms | 875 |
| | | | | | [H4NαacK16acK20me2] | 2 | 5.6 | 16.7 | 1-12, 98-102 | 34.7 | | |
| | | | | | [H4NαacK12me2K16ac] | 3 | 5.4 | 15.7 | 1-11, 98-102 | 33.7 | | |
